# Impact of the RaS-RiPP tryglysin and culturing conditions on *ex-vivo* oral microbiomes

**DOI:** 10.1101/2025.04.21.649776

**Authors:** Britta E. Rued, Achal Dhariwal, Brett C. Covington, Mohammad R. Seyedsayamdost, Sophie A. Krivograd, Russell P. Pesavento, Michael J. Federle, Fernanda C. Petersen

## Abstract

Natural products are ubiquitously produced by many species that we encounter during our daily lives. One genus, *Streptococcus*, can produce a wide array of quorum sensing linked natural products known as RaS-RiPPs (ribosomally synthesized and post-translationally modified peptides). Their production is triggered by the induction of an Rgg-SHP quorum sensing system, which senses the presence of SHPs (short hydrophobic peptides) and induces the gene expression of these operons. Previous work has found that streptococcal RaS-RiPPs modulate the growth of different streptococci and might play a role in antibiotic tolerance. This is of particular importance to the oral microbiome, where streptococci are a predominant genus. This study examines the impact of the RaS-RiPP Tryglysin A on *ex-vivo* oral systems and provides the groundwork for determining how producer Streptococci might impact the composition and function of oral microbiome communities. We additionally explore important factors to consider when culturing *ex-vivo* oral systems.

## INTRODUCTION

The oral cavity is a site of colonization for many important microbes, both bacterial and fungal^1–4^, which can have huge impacts on human health. These impacts include the initiation and development of dental caries, periodontal diseases, and enhanced development of oral cancers^5,6^. As such, understanding how microbial factors contribute to the establishment and maintenance of oral systems is of great importance, as they can impact the speciation and nature of the oral microbiome. A major constituent of the oral microbiome is the genus *Streptococcus*. Streptococci make up approximately 10-60% of the oral microbiome, depending on the niche examined and sequencing techniques used ^3,7,8^. Some of these organisms are keystone colonizers and possess the ability to drastically impact microbial balance^4,9^. Recently, we discovered that *Streptococcus mutans* and *Streptococcus ferus* can alter the growth of other oral streptococcal species via small post-translationally modified peptides called tryglysins ^10^. Tryglysins form a new class of post-translationally modified peptides known as “RaS-RiPPs”^11^, with **RaS** and **RiPP** standing for **Ra**dical **S**-adenosylmethionine enzyme and **Ri**bosomally translated and **p**ost-translationally modified **p**eptide, respectively^12^. To date, sixteen different classification groups of streptococcal RaS-RiPP peptides that form unique heterocyclic and macrocyclic structures have been described^11,13^. The tryglysin system in *S. mutans* is induced via quorum sensing and inhibits species such as *S. mitis, S. oralis*, and S*. sanguinis* at concentrations as low as 100 nM^10^. Owing to this potent activity, we wondered what impact these peptides might have on a complex oral system that includes multiple oral species.

Current models of complex oral systems have placed substantial efforts into optimizing culturing conditions. Most frequently used is SHI medium under anaerobic or microaerophilic conditions, replicating growth in a biofilm-like context^14–17^. SHI medium is a nutrient-rich and multi-component medium designed by Edlund *et al.,* 2013, to support the growth of a diverse oral community^14^. Edlund *et al*. and other investigators have shown that under anaerobic conditions saliva inoculation into SHI medium supports a diverse community^14,18^. These studies have demonstrated that culturing in SHI results in the growth of *Streptococcus*, as well as *Veillonella*, *Gemella*, *Granulicatella*, *Klebsiella*, *Lactobacillus*, and *Fusobacterium*^14^. Anaerobic conditions are primarily used due to the observation that many oral species are strict or obligate anaerobes, and previous studies found that saliva has a lower oxygen content^3,4,19^. Biofilms are typically considered to be more relevant to oral systems due to their role in the formation of plaque and establishing oral communities on the dental surface^4,20^. Few investigations have examined the impact of other types of growth conditions, be it the impacts of medium, saliva, oxygen or the type of growth model. In the development of experimental conditions to test tryglysin activity on oral microbes, we found that environmental settings were a critical variable.

The impact of culturing conditions on the fungal component of the microbiome is an additional consideration. Fungi are normal constituents of the oral microbiome and have significant effects on oral health^1,21^. However, fungi are less studied in oral systems due to their lower abundance, the difficulty of cultivation, lower sequencing database reliability, and the relative low rate of detection in shotgun metagenomics studies^1,2,21^. Fungal genera growth under anaerobic conditions has typically been observed to be low^16^.

Herein, we describe the first report of tryglysin activity on oral saliva samples. Our initial attempts to observe the impacts of tryglysin on oral microbiota were hampered by barriers associated with culturing conditions. We discovered that culturing in streptococcal chemically defined medium (CDM) was necessary for the activity of the RaS-RiPP tryglysin and that this medium selectively favored the growth of streptococci, even from a rich starting salivary inoculum. We also determined that the type of growth model used and the saliva and oxygen contents had substantial effects on the development of the oral consortia, and that serial culturing led to shifts in the detected species. Finally, we found that 5% CO_2_ allowed for the growth of fungal members of the oral community, making this a culturing method worth considering for future oral studies.

## RESULTS

### Tryglysin A inhibits the growth of saliva-derived oral species in a chemically defined medium that favors *S. salivarius*

Previous studies have demonstrated that tryglysins can inhibit the growth of many streptococcal species ^10^. As such, we reasoned that tryglysins should have the capacity to modulate the growth of oral streptococci and potentially the oral microbiome. To examine this aspect further, we first determined if tryglysin A (TryA) could inhibit the growth of sensitive streptococci in media typically used for *ex-vivo* culturing of oral species. One such medium that is frequently used in the oral microbiome field is SHI^14–16,22,23^, and a previous study demonstrated that other synthetic antimicrobial peptides (e.g., C16G2) were effective against streptococci in this medium^24^. To examine SHI, we exposed the TryA-sensitive species *S. mitis*^10^ to an inhibitory level of TryA or a peptide with no known function in quorum sensing or growth (reverse SHP or revSHP) in SHI medium (sequences listed in Table 1). As SHI medium contains blood and hemin, their inherent turbidity and light-absorbing properties are confounders in optical measurements. As such, viability of microbes was determined by plating for CFU/mL over time (Fig. 1A). To our surprise, SHI medium abrogated any significant inhibition of TryA against *S. mitis* (Fig. 1A). This was not due to the absence of tryglysin activity, as this same batch of TryA was effective against *S. mitis* in a chemically defined medium (CDM, Fig. 1B). This was also not due to a difference in lab-specific strain backgrounds, as retesting with the same strain of *S. mitis* in a separate laboratory setting produced the same results: unobservable activity in SHI medium (Fig. S1A). We proceeded by culturing a previously collected saliva inoculum^16^ in CDM anaerobically^14,15^, since *S. mitis* inhibition was reproducibly observed in this medium. Experimental conditions are outlined in Table 2 and subject demographics for this saliva collection, designated “Norway” are listed in Table 3. We first determined if inhibition was apparent by growing salivary inoculum in CDM, as well as by measuring pH and optical density over time. The salivary inoculum was placed in CDM and exposed to a range of TryA concentrations, revSHP, or PBS. We observed that TryA delayed growth of the salivary inoculum in a dose dependent manner and this was reflected in the delayed acidification of the medium (Fig. 1C-1D), whereas revSHP and PBS treatments grew rapidly (Fig. 1C-1D), consistent with previous observations for mono-species cultures of streptococci ^10^. CFU/mL was not drastically reduced over time (Fig. 1E, Fig. S1C). We performed 16S rRNA sequencing on these samples after 24 hours and primarily observed streptococci (Fig. S1B). To achieve species-level resolution, we carried out shotgun metagenomics and analyzed the data with MetaPhlAn3. From this analysis, we found a high predominance of *Streptococcus salivarius* under all conditions, indicating strong selection for this species after 24 hours (Fig. 1F, Supplementary File 2: Table 1). Other streptococcal species were observed, but their abundances were low (less than 0.2-2% relative abundance) and varied between the sample conditions (Fig. 1F, Supplementary File 2: Table 1). These observed differences were not due to the initial salivary inoculum being skewed in composition, as direct sequencing of saliva showed a diverse composition of phyla and genera of both bacteria and fungi (Fig. 2A, 2C, Fig. S1E). Differential abundance analysis using DESeq2^25^ revealed that TryA treated samples had significantly higher levels of *Candidatus Saccharibacteria* than both PBS and revSHP treated samples (Adjusted P-value < 0.05; Supplementary File 2: Table 2). *S. parasanguinis* levels were sensitive to the addition of either TryA or revSHP (Supplementary File 2: Tables 1-2). No significant differences between samples were observed via alpha diversity metrics such as Chao1 and Shannon (Supplementary File 2: Table 3). Principle component analysis (PCoA) analysis of MetaPhlAn3 results showed that TryA treated samples clustered away from other treatment conditions, but these differences were non-significant by PERMANOVA (R^2^ = 0.403, P-value = 0.467) (Fig. S1D). We conclude that CDM leads to strong selection for streptococci under anaerobic conditions, and that TryA addition results in additional selective pressure. It also suggests that TryA may have effects on the presence of oral bacteria such as *Candidatus Saccharibacteria*, organisms that form obligate relationships with other bacteria for their survival^26^. Even though these results were limited due to the collapse of the consortia to streptococci, the findings demonstrate an advantage over single laboratory isolate studies. Typically, single-species models are limited to using “domesticated” laboratory strains that have been propagated in culture for years, potentially altering their behavior compared to their *in vivo* counterparts. Therefore, a model that predominantly consists of streptococci from human saliva could be particularly relevant for understanding species-specific interactions in streptococcal communities. CDM culturing of saliva represents one such model that can be used in this respect. In addition to this, these findings represent the first examination of the impact that a streptococcal RaS-RiPP might have on oral microbiomes.

**FIGURE 1:**
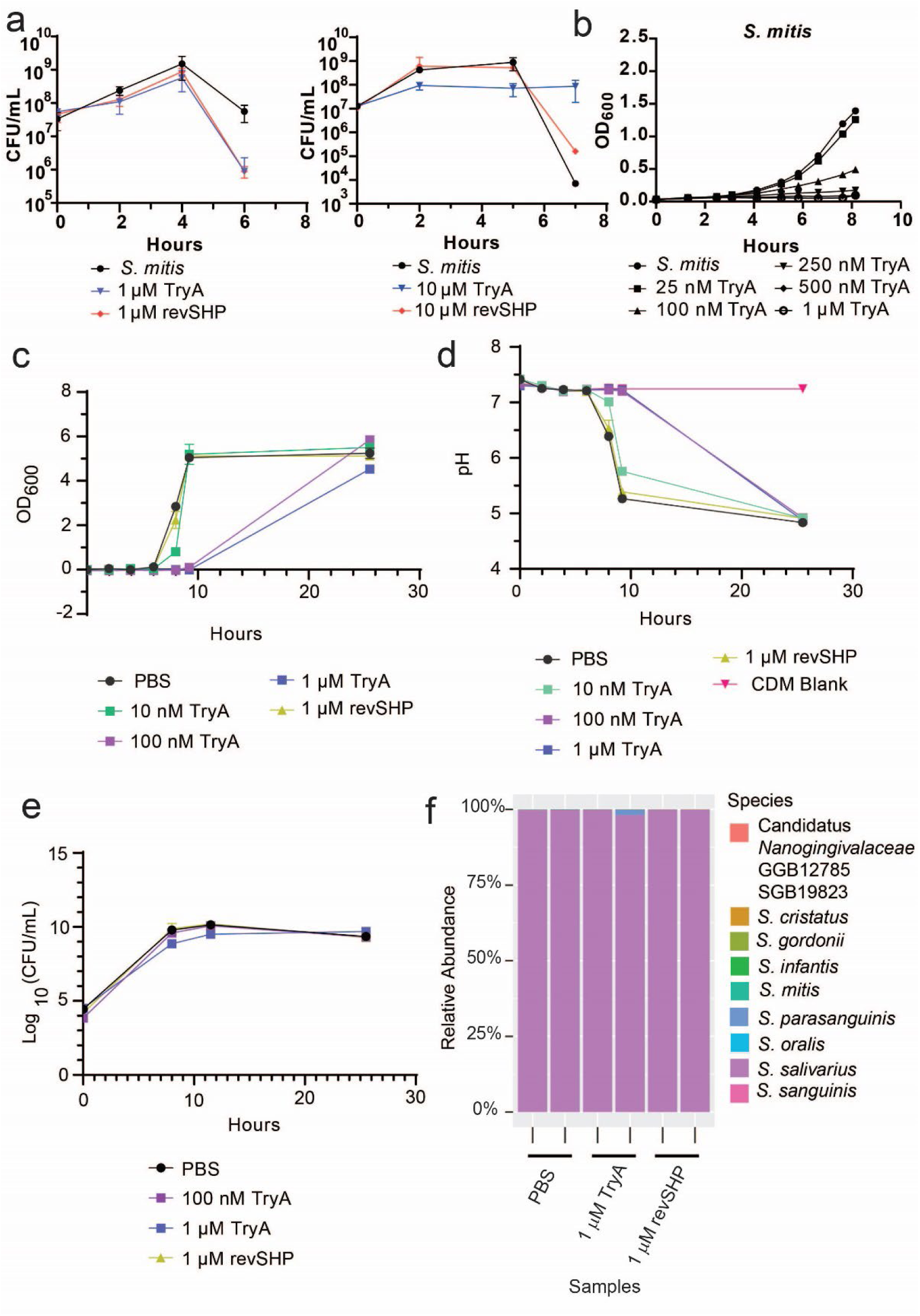
Examination of Tryglysin A’s effectiveness in CDM vs. SHI media, and response of salivary inoculum in chemically defined media (CDM) to the addition of Tryglysin A (TryA), a control peptide that does not affect growth or quorum sensing (reverse SHP/revSHP), or PBS. Concentrations of TryA are indicated in the legend below graphs in A through E and correspond to the respective symbol types. A) CFU/mL of wild-type *S. mitis* exposed to increasing TryA, in SHI media. Concentrations of TryA added are indicated below the graph. This experiment was performed twice: once with 1 µM TryA, once with 10 µM TryA. Graphs for both experiments are shown. B) Growth curve of wild-type *S. mitis* exposed to increasing TryA in CDM. This experiment was performed three times with similar results. C) Growth curve of salivary inoculum in CDM over time. Concentrations of TryA added or other conditions are indicated below the graph. This experiment was performed three times with similar results. D) Examination of pH of salivary inoculum in CDM over time. Concentrations of TryA added or other conditions are indicated below the graph. This experiment was performed four times with similar results. E) Separate experiment examining CFU/mL of salivary inoculum in CDM over time. Concentrations of TryA added or other conditions are indicated below the graph. This experiment was performed twice and correlates with data shown in Fig. S1C. F) Speciation as determined via shotgun metagenomics for samples grown in the presence of PBS, 1 µM TryA, or 1 µM reverse SHP peptide in CDM for 24 hours. Relative abundance is plotted for each sample. Species detected are indicated by the legend below the graph.

**FIGURE 2:**
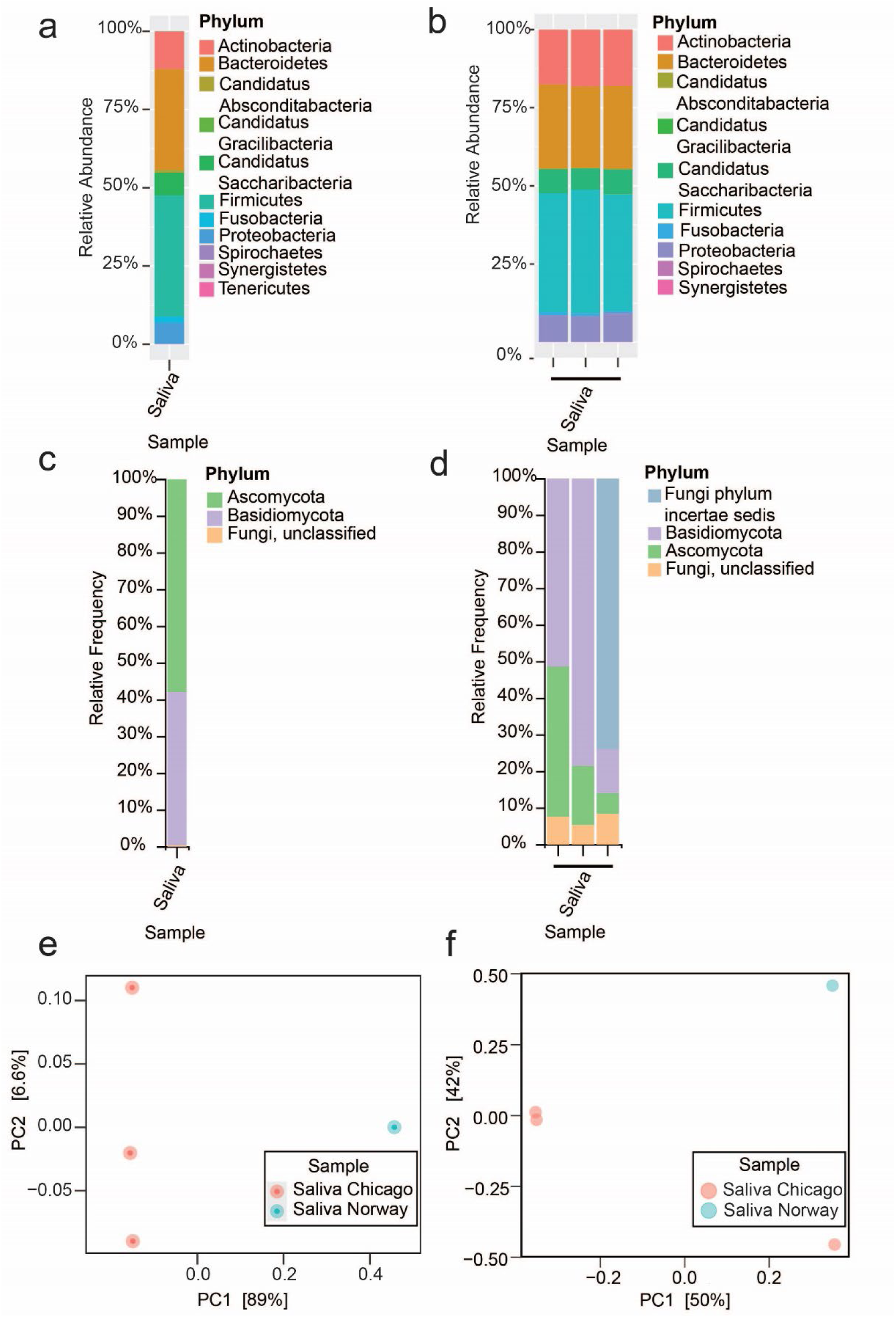
Comparison of the bacterial and fungal phyla detected in human pooled saliva isolates. Relative abundance is shown for each sample, phyla are indicated by color as detailed in the legends beside each graph. The Norway pooled saliva was sequenced once, whereas the Chicago pooled saliva was sequenced in triplicate, with all samples originating from DNA extracted from the same salivary pool. A) Bacterial phyla from shotgun metagenomics analysis of pooled human saliva from Norway. B) Bacterial phyla from shotgun metagenomics analysis of pooled human saliva from Chicago. Note that this data is also presented alongside additional samples in Fig. 3A. C) Fungal phyla detected in pooled human saliva from Norway by ITS sequencing. D) Fungal phyla isolates detected in pooled human saliva from Chicago by ITS sequencing. Note that this data is also presented alongside additional samples in Fig. 3B. E) Jaccard PCoA of Chicago and Norway pooled saliva from shotgun metagenomics, at the species level. F) Bray-Curtis PCoA of Chicago and Norway pooled saliva from ITS sequencing.

**TABLE 1.** Amino acid sequence of peptides used in this publication.^a^.

| Peptide | Amino acid sequence |
| --- | --- |
| *S. ferus* tryglysin A | VNSWGKH, modification at WGK |
| revSHP peptide, *S. mutans* reverse SHP | GGGIIIITE |

<sup>a</sup>*S. ferus* tryglysin A was purified directly from *S. ferus* culture supernatants, whereas *S. mutans* reverse SHP was synthesized by ABClonal. See *Materials and Methods* for further details.

**TABLE 2.** Experimental parameters used in this study to examine impact of culturing conditions on *ex-vivo* oral microbiomes.

| Goal | Number of replicates per experimental condition | Ex-vivo saliva sources | Incubation conditions before sequencing | Community sequencing technique used<sup>a</sup> |
| --- | --- | --- | --- | --- |
| Examine impact of tryglysin A on *ex-vivo* oral microbiomes. | 2 | Norway | Anaerobic, glucose CDM<sup>b</sup>, 24 hours, planktonic<sup>d</sup> | 16S sequencing, shotgun metagenomics |
| Determine impact of pooled human saliva source on *ex-vivo* speciation. | 1 for Norway, 3 for Chicago | Norway, Chicago | No culturing, direct extraction | ITS sequencing, shotgun metagenomics |
| Examine impact of biofilm vs. planktonic culturing methods. | 3 | Chicago | 5% CO<sub>2</sub>, glucose SHI<sup>c</sup>, 24 hours, planktonic or biofilm<sup>d</sup> | 16S sequencing, ITS sequencing |
| Examine impact of CDM vs. SHI medium. | 3 | Chicago | 5% CO<sub>2</sub>, glucose SHI or CDM<sup>c</sup>, 24 hours, planktonic<sup>d</sup> | 16S sequencing, ITS sequencing |

<sup>a</sup>16S sequencing, ITS sequencing, and shotgun metagenomics sequencing were performed as detailed in *Materials and Methods*.
<sup>b</sup>CDM stands for streptococcal chemically defined medium.
<sup>c</sup>SHI stands for SHI medium, from Edlund *et al.*, 2013. *Microbiome*.
<sup>d</sup>Planktonic cultures were grown in broth suspension in 15 mL tubes, whereas biofilm cultures were grown as 1 mL cultures in 24 well plates. For further details see *Materials and Methods*.

**TABLE 3.** Subject demographics for saliva collection at University of Illinois - Chicago^a^ and University of Oslo^b^.

| Demographic | Chicago<br>N | Chicago<br>Percent | Norway<br>N | Norway<br>Percent |
| --- | --- | --- | --- | --- |
| **Age group** |  |  |  |  |
| 18-24 | 1 | 12.5% | 0 | 0%<sup>c</sup> |
| 25-34 | 5 | 62.5% | 0 | 0%<sup>c</sup> |
| 35-44 | 1 | 12.5% | 7 | 87.5% |
| 45-54 | 1 | 12.5% | 0 | 0%<sup>c</sup> |
| >55 | 0 | 0%<sup>c</sup> | 1 | 12.5% |
| **Gender** |  |  |  |  |
| Male | 5 | 62.5% | 2 | 25% |
| Female | 3 | 37.5% | 6 | 75% |

<sup>a</sup>All data collected at University of Illinois – Chicago for this study was collected at the University of Illinois Chicago CCTS Clinical Research Center, under UIC Research IRB Study ID 2023-0288.
<sup>b</sup>All data collected at University of Oslo for this study was collected at University of Oslo under Norwegian Regional Ethics Committee Study ID REK20152491.
<sup>c</sup>0% indicates no samples were collected from this age group category

### Collection of a secondary salivary pool and effects of growth model and oxygen presence

Due to our findings regarding CDM growth of salivary inocula, we wondered if this streptococcal selection was reproducible and if we could identify conditions that would allow for more diverse speciation for our assays with CDM or other media. Thus, we collected a secondary saliva pool at the University of Illinois Chicago and determined how parameters such as media composition, presence of CO_2_ and oxygen, initial starting saliva proportions, and growth model affected speciation outcome (Table 2). Subject demographics comparing saliva collections from Chicago and Norway are shown in Table 3. Saliva samples were collected, pooled, and grown under various conditions.

The species composition of the pooled saliva collection from Chicago was compared to the Norway salivary collection. Overall, bacterial phyla and genera detected by shotgun metagenomics were similar between the two collections, aside from the low but measurable presence of an uncharacterized mycoplasma species of *Tenericutes* in the Norway collection not present in the Chicago collection (Fig. 2A-2B, Supplementary File 2: Tables 4-5). Variation between the two collections was present but non-significant, as observed via Jaccard PCoA and PERMANOVA (R^2^ = 0.957, P-value = 0.25) (Fig 2E). We also examined fungal composition by ITS sequencing. This demonstrated that the previous saliva collection was composed primarily of *Ascomycota* and *Basidiomycota*, with the genus *Malassezia* being dominant (Fig. 2C, Fig. S1E, Supplementary File 2: Table 6). Composition of collections were similar at the phylum level but differences were observed in fungal genera (Fig. 2C-2D, Fig. S1E-S1F). For instance, the Norway collection contained *Penicillium* which was not observed in the Chicago collection, whereas the Chicago collection contained Cryptococcus that was unobserved in the Norway collection (Fig. S1E-S1F, Supplementary File 2: Tables 6-7). Further analysis found no significant differences by alpha diversity (Simpson’s Dominance, Shannon Entropy, Chao1 Index and Faith’s) or beta diversity metrics (PERMANOVA: R^2^=0.532, P-value = 0.25) (Supplementary File 2: Table 8). Bray-Curtis PCoA also reflects these findings (PERMANOVA: R^2^= 0.446, P-value = 0.5; Fig. 2F). However, this interpretation also relies on the fact that the Norway consortia was only sequenced once due to the limited availability of biomaterial.

We next decided to look at the impact of growth models (biofilm vs planktonic) and oxygen content on bacterial speciation. Culturing techniques used are outlined in Table 2. Chicago salivary samples were grown under 5% CO_2_ and compared to the uncultured, pooled Chicago saliva sequencing via shotgun metagenomics, 16S and ITS sequencing, as well as previously published results^14^. We found that the relative frequency of bacterial and fungal phyla decreased in both biofilm and planktonic samples compared to saliva samples (Fig. 3A-3B). Both biofilm and planktonic samples were predominated by Firmicutes for bacteria and Ascomycota for fungi, which correlated with *Streptococcus*, *Cyberlindera*, and *Candida* levels respectively (Fig. 3A-3B, Fig. S2A-S2B). Bacterial and fungal genera were detected and a summary of the ASV counts from QIIME2 analysis and relative abundance of shotgun metagenomics analysis are listed in Supplementary File 2: Tables 5, 7 and 9. Alpha diversity metrics for biofilm and planktonic samples were significantly different for 16S sequencing results (Fig. S2C), but not ITS sequencing (Fig. S2D). Beta diversity demonstrated that for both 16S and ITS sequencing, compositions of samples were distinct (Fig. S3A-S3B). PCoA and PERMANOVA indicated samples were significantly different from each other (R^2^=0.993, P-value = 0.002 for shotgun metagenomics; R^2^=0.449, P-value = 0.018 for ITS) (Fig. 3C-3D). We note that these biofilm samples were not grown in plates with a pre-treated saliva coating, which marks one additional change besides the culturing in 5% CO_2_ compared to previous culturing studies^14–16^. However, we find that like the previous Edlund *et al.,* 2013 study using the pre-coated saliva method, streptococci are predominant, although the relative percentage of streptococci increases under 5% CO_2_ without a saliva coat (Fig. S2A, Supplementary File 2: Table 10. In planktonic samples, streptococci make up 81.8% of genera detected (Fig. S2A, Supplementary File 2: Table 10). *Veillonella, Gemella, Granulicatella, Lactobacillus, Prevotella, Porphyromonas, Neisseria* were also detected in our study, though at lower proportions than reported in Edlund *et al.* More bacterial genera were detected in the planktonic samples in our study than in either biofilm culturing methods (Supplementary File 2: Table 10). Additionally, we observed that samples cultured at 5% CO_2_ had observable levels of fungi. Previous studies have observed fungal levels using a pre-coated saliva biofilm model with SHI medium to be low during anaerobic culturing^16^. In contrast, our samples were dominated by detectable *Cyberlindnera* during biofilm culturing and *Cyberlindnera* and *Candida* during planktonic culturing (Fig. 3B, Fig. S2B, Supplementary File 2: Table 7).

**FIGURE 3:**
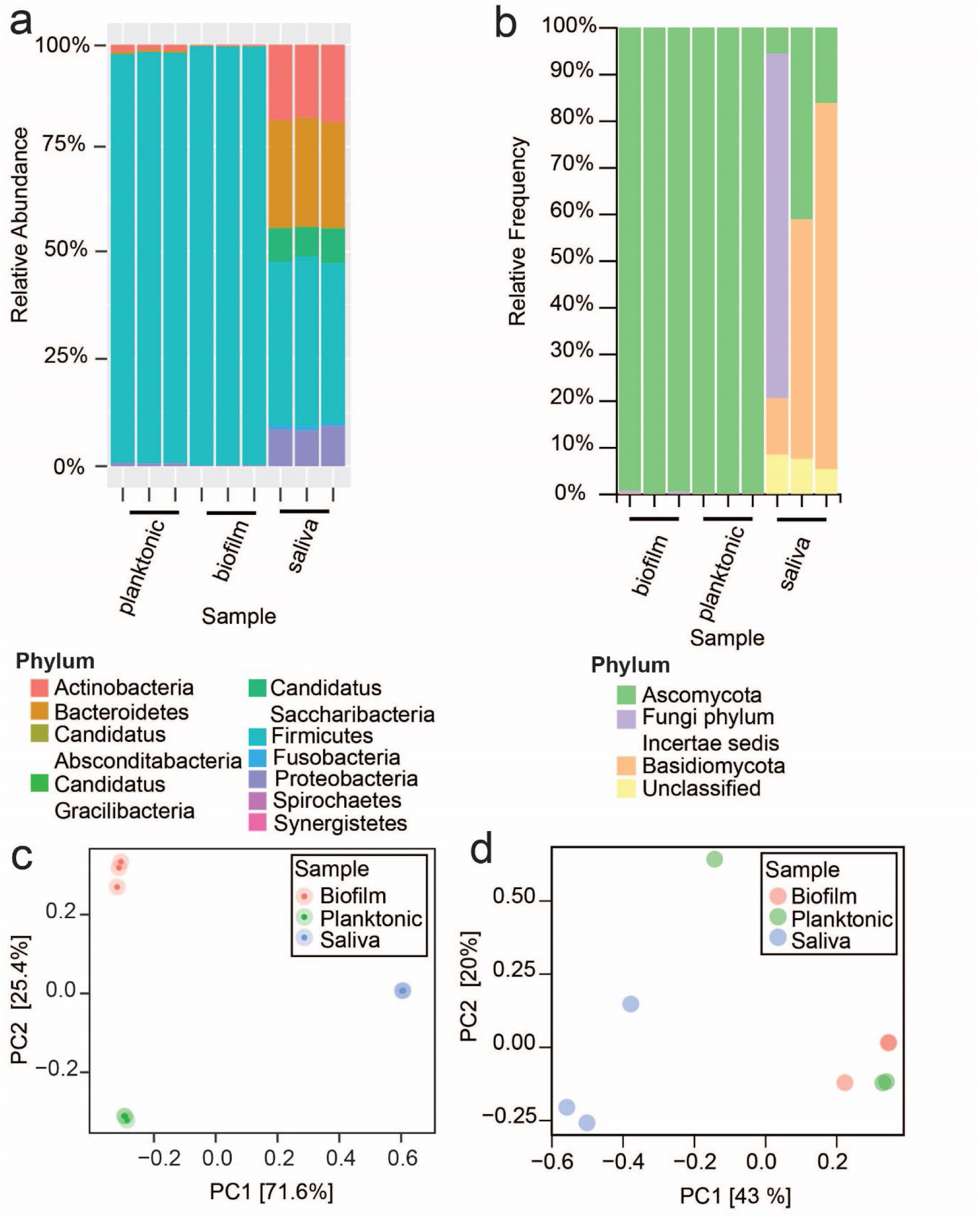
Examining the use of 5% CO_2_ as a culturing method for *ex-vivo* oral microbiomes. All samples shown were grown in SHI media for 24 hours with 5% CO_2_. A) Shotgun metagenomics results showing bacterial phyla detected in a biofilm or planktonic model of growth, compared to phyla detected in the original salivary inoculum. B) ITS sequencing results showing fungal phyla detected in a biofilm of planktonic model of growth, compared to phyla detected in the original salivary inoculum. C) Jaccard PCoA of samples in panel A from shotgun metagenomics sequencing at the species level. D) Bray-Curtis PCoA of samples in panel B from ITS sequencing.

Overall, these data indicate that although not perfect, culturing saliva consortia with 5% CO_2_ recapitulates the presence of major oral genera for bacteria and allows the propagation of certain fungi. This represents a much more accessible method of culturing oral microbiota than prior methods, as it does not necessitate an anaerobic chamber. It also indicates that this method might provide a means to assess oral fungal-bacterial interactions.

### Impact of media conditions and re-culturing on consortia results

We determined that culturing the Norway salivary sample in CDM led to consortia collapse to streptococci (Fig. 1F), To further examine this finding, we tested whether selection would be similar with the Chicago consortia using the 5% CO_2_ planktonic model with CDM or SHI medium, as well as to examine the impact of serial culturing on species diversity. SHI medium cultures grown for 24 hours were frozen at -80°C as glycerol stocks and then thawed, split, and used to inoculate fresh SHI or CDM media as triplicate samples for 24 hours and then examined for their bacterial and fungal composition. Total CFU/mL displayed no differences between serially cultured samples or cultured saliva. In CDM, serial culturing in SHI medium resulted in a higher overall CFU/mL (Fig. 4A). No major differences in gross colony morphologies were observed between serially cultured samples and samples directly inoculated with saliva in either medium (Fig. 4B).

**FIGURE 4:**
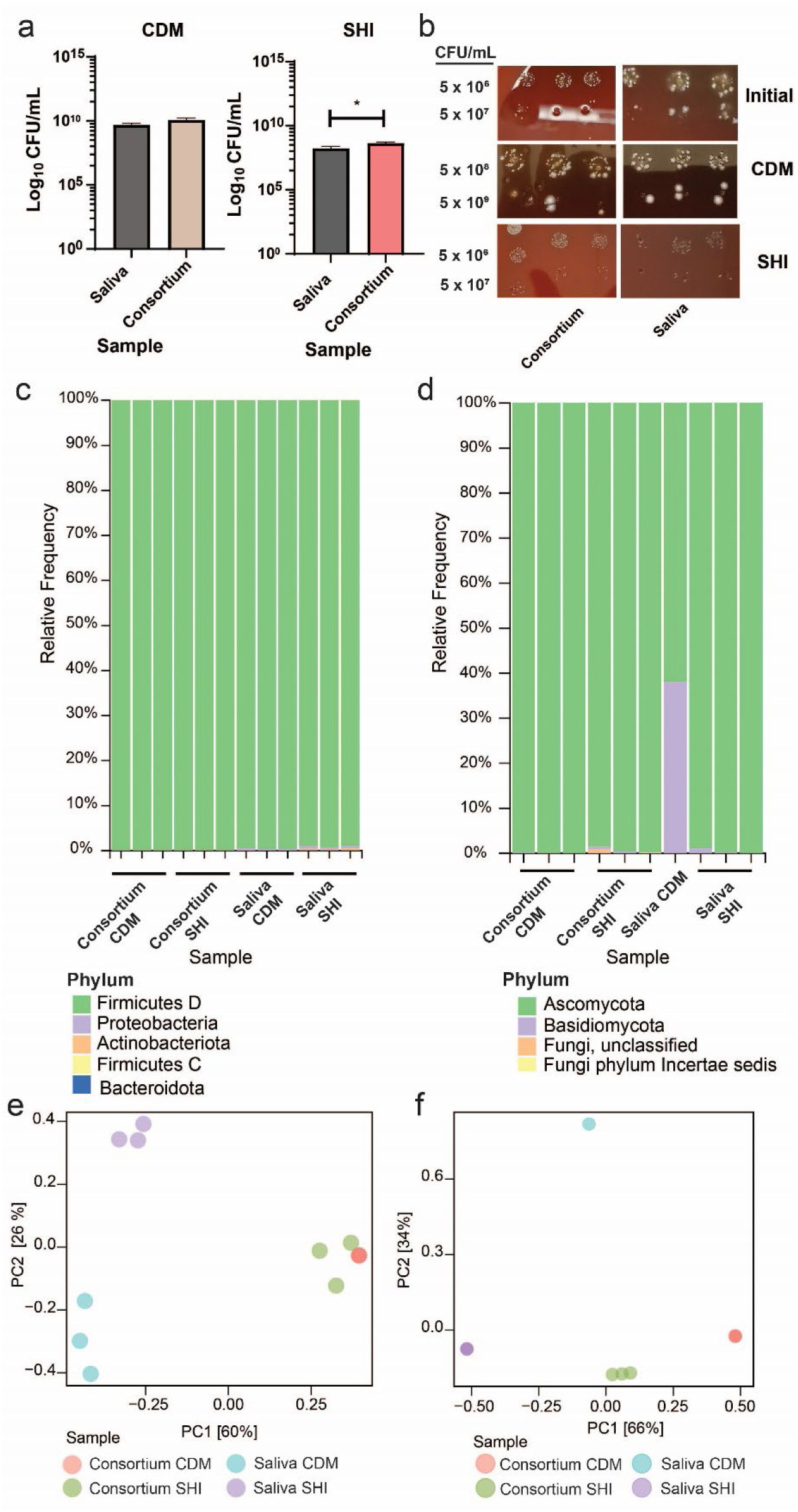
Examining the impact of CDM vs. SHI media for culturing *ex-vivo* oral microbiomes. Samples were grown in CDM or SHI media for 24 hours with 5% CO_2_, in biological triplicate. A) CFU/mL of saliva inoculum or previously stored consortia after growth in CDM or SHI medium. These experiments were performed twice with similar results. B) Colony morphologies observed for saliva or consortium propagation in CDM or SHI media after 24 hours of growth. Shown also are the initial colony morphologies observed in diluted saliva or consortium. This experiment was performed twice with similar results. C) 16S sequencing results showing bacterial phyla detected after growth of saliva inoculum or previously stored consortia in CDM or SHI media. D) ITS sequencing results showing fungal phyla detected after growth of saliva inoculum or previously stored consortia in CDM or SHI media. E) Jaccard PcoA of samples in panel C from 16S sequencing. F) Bray-Curtis PcoA of samples in panel D from ITS sequencing.

We then examined bacterial composition by sequencing and made several important findings. We observed that the use of serially cultured samples selects for specific bacterial genera in both CDM and SHI medium: *Streptococcus* and *Ligilactobacillus* (Supplementary File 02: Table 11, Fig. 4C, Fig. S4A). The relative amounts of these change depending on the medium condition, with SHI selecting for more *Ligilactobacillus* and CDM selecting for more *Streptococcus* (Fig. S4A; Supplementary File 02: Table 11). In contrast, direct culturing of saliva in SHI medium leads to a higher number of bacterial genera detected than in CDM (total of 13 vs. 8). We also found that culturing saliva directly in CDM led to the growth of primarily streptococci (Fig. S4A; Supplementary File 02: Table 11), similar to the results from our initial investigations (Fig. 1F). These differences are reflected in the alpha and beta diversity analysis of these results (PERMANOVA: R^2^=0.876, P-value = 0.001) (Fig. 4E, Fig. S4B-S4C, S5). For alpha diversity, Simpson’s Dominance and Shannon Entropy were statistically significant between CDM and SHI medium, with SHI medium having higher relative values for these metrics (Fig. S5A) indicating that species richness, relative abundance, and relative evenness are higher for samples cultured in SHI medium. In contrast, Faith’s Phylogenetic Diversity and Chao1 Index were only significant when comparing the use of inoculum source (Fig. S5A). This indicates that genera richness decreases when using re-cultured consortium as a source. Finally, beta diversity examined using Jaccard distance, and was significantly different when comparing inoculum sources only, indicating that this condition is key for determining the number of unique species (PERMANOVA: R^2^=0.572, P-value = 0.003) (Fig. S4C, S5B). This heavily suggests that continually re-cultured samples in SHI medium should be used with caution for oral microbiome studies, as a clear skew in genera representation results.

We then compared fungal composition from the same samples. Similar to previous results for planktonic culturing in SHI medium, *Ascomycota* was the dominant phylum (Fig. 3B, Fig. 4D, Supplemental File 02: Table 12). Continued culturing in either medium led to predominance of *Candida* (Fig. S6A, Supplemental File 02: Table 12), indicating that re-culturing leads to a selection for specific fungal genera. No pairwise differences in alpha or beta diversity via Bray-Curtis were observed between samples (Fig. S6B-S6C). Overall microbiome composition was significantly different between groups (PERMANOVA, R^2^= 0.997, P-value = 0.001) (Fig. 4F, Fig. S6C). Significant differences were observed in alpha and beta diversity when examining medium conditions alone (PERMANOVA, R^2^ = 0.543, P-value = 0.005) (Fig. S7A-S7B). Inoculum source was also significant when compared alone (PERMANOVA, R^2^ = 0.449 P-value = 0.015) (Fig. S7B). This indicates that the differences in re-culturing and media composition of fungi are significant for fungal species diversity. However, this analysis could also be affected by the lower number of fungal reads obtained for this experiment and should be interpreted with some caution.

## DISCUSSION

Our initial investigations started by examining the impact of tryglysin A on the growth of an *ex-vivo* oral system. Initially, we found that TryA was ineffective in SHI medium (Fig. 1A), prompting us to switch to culturing in CDM, a peptide free medium known to support TryA activity^10^ (Fig. 1B). This phenomenon, where peptides show activity in minimal but not rich media, has been observed previously^27–29^. In CDM, TryA concentrations ranging from 100 nM to 1 µM inhibited the salivary inoculum (Fig. 1C-E, Fig. S1C), although it led to a notable selection for streptococci across all samples (**Fig. 1F**). Despite this selective limitation, we observed that TryA significantly affected the presence of *S. parasanguinis* and a member of Candidatus *Saccharibacteria*, suggesting that TryA can modulate certain members of the oral microbiome (Fig. 1, Supplementary File 02: Table 2). This represents the first observation of streptococcal RaS-RiPPs influencing their residing oral systems.

In the second part of our study, we aimed to further explore conditions that could influence outcomes in *ex-vivo* models of the oral microbiota. Previous studies of oral consortia have primarily been performed in anaerobic chambers with atmospheres of varying mixtures of carbon dioxide, nitrogen, and hydrogen^14–16,22^. These conditions allow for the excellent growth of strict anaerobes found in the oral microbiome. However, this does not mean that other culturing methods should not be explored for their efficacy, as other studies have supported the idea that an oxygen gradient occurs in the oral microbiome^3,30^. Saliva has been traditionally considered a low oxygen environment due prior reports of salivary oxygen levels around 0.08 ppm and rapid turnover of oxygen by oral organisms^19,31^. However, a new study by Podunavac *et al.,* found drastically varying levels of oxygen and carbon dioxide between conditions and human volunteers^32^, although this could be partially attributed to sampling methods. Additionally, data from Sikdar *et al.* 2024 supports the idea that 5% CO_2_ is a viable culturing method for *ex-vivo* oral systems^33^. In their study, they found that a supragingival plaque system only produced n-acyl homoserine lactone signaling molecules under the presence of 5% CO_2_, and that these significantly impacted the oral system^33^. Also similar to our findings, they found that under 5% CO_2_ in both a planktonic and biofilm type model *Streptococcus* predominated^33^. This study together with our data indicates that 5% CO_2_ should be considered as a secondary culturing method for oral samples. This additionally makes culturing oral microbiomes more accessible to other laboratories, as anaerobic culturing requires an expensive investment in equipment that limits oral microbiome experiments to those who possess these set-ups.

Culturing in 5% CO_2_ had a clear impact on bacterial phyla and speciation, strongly selecting for Firmicutes (Fig. 3A), but also allowing the growth of fungal species (Fig. 3B). Anaerobic culturing of *ex-vivo* systems have been reported to have low fungal presence^16^ but we observe that during 5% CO_2_ culturing in SHI medium fungi are present (Fig. 3B, Fig S2B). This is not dependent on the growth model used, as we observed fungi were present in both a biofilm and planktonic model (Fig. 3B, Fig. S2B). This presence of fungi was dependent on the use of SHI medium, as the use of CDM resulted in fewer fungal ASVs by ITS, and some samples generated few reads (Fig. 4D, Supplementary File 02: Table 12). Re-culturing of oral inocula appeared to select for the presence of specific genera of fungi such as *Cyberlindnera* and *Candida* (Fig. S6A). We assert that due to these data and findings concerning other important factors such as AHL production^33^, this culturing method should be used to examine additional aspects of oral microbiome interactions.

In our culturing experiments, we observed differences between growth models at the bacterial and fungal genera level (biofilm vs. planktonic) (Fig. S2). In our initial experiments, a larger salivary inoculum was used for planktonic samples (Fig. S2). As such, the difference between higher saliva concentration and mode of growth cannot be separated. One thing it does emphasize is the fact that both saliva concentration and mode of growth contribute to *ex-vivo* speciation outcome. This is not surprising, as other studies have found human saliva can greatly alter growth and behaviors of oral species^34,35^. We also observed clear impacts of re-culturing samples from SHI medium. Originally, we hypothesized that SHI could lead to stable and re-culturable consortia for use in the laboratory. However, we found that there were clear impacts at the genus level for both bacteria and fungi for re-culturing (Fig. S4, Fig. S6). Re-culturing might be a viable method for selecting for simpler fungal and bacterial consortia, but it clearly results in certain biases.

From this study, we conclude that multiple factors must be carefully considered in *ex-vivo* oral culturing studies, including the role of atmosphere, culturing method, initial inoculum, and appropriate medium conditions. Optimizing these aspects is essential for developing robust *ex-vivo* oral models to better understand physiological aspects of the oral microbiome. Notably, while CDM supported fewer species compared to the rich SHI medium traditionally used, it provided valuable insights into the activity of TryA, which was ineffective in SHI. This highlights the importance of selecting the appropriate medium to achieve specific peptide activity or antimicrobial agents in general. The phenomenon wherein certain peptides are functional in a defined medium but not in a rich medium remains an intriguing question. In the meantime, it is important to explore diverse conditions for studying various compounds that may exhibit activity under different circumstances, thereby expanding our understanding and potentially uncovering new therapeutic applications.

## MATERIALS AND METHODS

### Bacterial strains and growth conditions for strains and *ex-vivo* oral samples

Bacterial strains used in this study are listed in Supplementary Data File 1: Table S1. *S. mitis* strains were derived from CCUG31611. Strains and *ex-vivo* oral microbiota samples from saliva were grown on Todd-Hewitt plates (TH, BD Biosciences) with 1.4% Bacto-Agar (BD Biosciences) and 0.2% Yeast Extract (VWR, RPI International), Tryptic Soy plates (TS, BD Biosciences) with 1.4% Bacto-Agar (BD Biosciences), Tryptic Soy plates with 1.4% Bacto-Agar (BD Biosciences) and 5% defibrinated sheep blood (QuadFive or Fisher Scientific), in Tryptic Soy Broth (TSB, BD Biosciences), in Todd-Hewitt (TH, BD Biosciences) with 0.2% Yeast Extract (VWR, RPI International), in SHI medium, or in streptococcal chemically-defined medium (CDM) plus 1% glucose at 37°C with 5% CO_2_.

The components and recipe for streptococcal CDM used were as described previously^36^. The components and recipe for SHI medium used were as described previously^14,16^.

### Purification of tryglysin A peptides and synthesis of *S. mutans* reverse SHP peptides

*S. mutans* reverse SHP peptide was purchased from ABClonal (Woburn, MA. Purities of peptide preparations used in assays were greater than 70%. *S. mutans* reverse SHP peptide was reconstituted as 10 mM stocks in DMSO and was stored in aliquots at -20°C. For experiments with *S. mutans* reverse SHP peptide subsequent dilutions for working stocks (1 mM, 100 µM, 10 µM) were made in DMSO or PBS and stored at -20°C. For sequences of peptides see Table 1.

For purification of tryglysin, *S. ferus* DSM 20646 was grown overnight in 10 mL of Todd Hewitt broth +0.2% yeast extract (5% CO_2_, 37° C) for 12-15 hours. The overnight culture was then centrifuged for 10 min at 4000*xg*, and the cell pellet was resuspended in 10 mL of CDM. The resuspended bacterial pellet was used to inoculate one liter of CDM in a glass bottle. This re-suspended culture was then incubated for 20–24 h at 37° C with 5% CO_2_. The resulting culture was centrifuged for 10 min at 4000*xg*, and cell-free supernatant filtered through 0.2 µm pore filters (VWR). 500 mL of the resulting filtered supernatant was loaded onto a 1g HyperSep Hypercarb SPE cartridge (Thermo Scientific, 60106-404), which had been preconditioned with 5–10 column volumes of 100% acetonitrile (ACN) with 0.1% formic acid (FA) followed by 5–10 column volumes of 100% H_2_O + 0.1% FA. The column was then washed with five column volumes of 100% H_2_O + 0.1% FA. Tryglysin A was eluted with 10–20 column volumes of 50% ACN + 0.1% FA. The eluted material was then evaporated to dryness and resuspended in 1 mL of H_2_O + 0.1% FA. Tryglysin A was purified further through two rounds of HPLC using an Agilent 1260 infinity system. In the first round, the resuspended samples were run on a semi-preparative scale with 100 μL injection volumes onto a Phenomenex Luna Omega Polar C18 100Å column (5 μm, 250 x 10 mm). Compounds were eluted using an initial 3.5-minute isocratic step running 100% H_2_O + 0.1% FA followed by a ramp to 10% ACN + 0.1% FA for 5 min, and a final ramp to 100% ACN + 0.1% FA over 2 min. During this time, 0.5 min fractions were collected. Fractions containing tryglysin A were identified through high-resolution QTOF-MS analysis using an Agilent 6540 UHD mass spectrometer. Fractions containing tryglysin were further purified through analytical HPLC with a Phenomenex Luna Omega Polar C18 100Å (1.6 μm, 150 x 2.1 mm) column using the same method. Several liters of culture were worked up in this manner to obtain sufficient tryglysin yields for experiments.

### Collection of salivary inoculums

For saliva samples used at the University of Oslo, samples (also designated ‘Norway’ collection) were collected as outlined in Dornelas-Figueria *et al.,* 2013^16^. These samples were collected in accordance with the Declaration of Helsinki and approved by the Norwegian Regional Ethics Committee (REK20152491) for studies involving human samples. For saliva samples used at the University of Illinois Chicago, samples (also designated ‘Chicago’ collection) were collected in conjunction with the CCTS Clinical Resources Center at the University of Illinois Chicago, in accordance with the Belmont Report and DHHS regulations 45 CFR Part 46 and approved by the University of Illinois Chicago Institutional Review Board (STUDY2023-0288). Sample collection, consent, demographic data collection, and de-identification prior to use by researchers for this study was conducted by CCTS Clinical Resources Center. Subjects were those over the age of 18, not currently using tobacco products, not under treatment with antibiotics or systemic disease, not recently diagnosed or having active COVID-19 infection, considered in good general health, and having good oral health (i.e. not currently having gum disease, excessive cavities, and possessing at least 20 teeth, in other words, not having undergone multiple extractions of teeth). Conglomerate demographics for subjects are listed in Table 3. Subjects were asked to expectorate 5 mL of saliva directly into a collection tube, after which samples were stored at 4°C. The same day of collection, samples collected and centrifuged at 2,600*xg* for 10 min to spin down large debris and eukaryotic cells, and resulting saliva supernatants were pooled together. Pooled saliva was stored with 20% glycerol at -70 °C, until all samples had been collected, at which time they were pooled again and then aliquoted into 1 mL volumes, which was referred to as pooled saliva. Pooled saliva was stored at -70 °C for use in experiments.

### Growth curves and CFU/mL assays of *S. mitis* exposed to tryglysin A

*S. mitis* pre-stored stocks with 20% glycerol at -70°C or -80°C were thawed and centrifuged at 10,000*xg* for 5 min, and then washed with 1 mL of CDM or SHI medium. Cultures were then resuspended in 1 mL of either CDM or SHI medium to be used for the experiment and diluted with according medium to required volume to OD_600_ ∼0.03. For growth curves in CDM, at 1 hour post inoculation (approximately OD_600_ ∼0.05-0.1), 25 nM, 100 nM, 250 nM, 500 nM, or 1 µM tryglysin A were added to cultures. Samples were grown at 37°C with 5% CO_2_, with OD_600_ observed every 45 min to 1 hour with a GENESYS 30 Vis spectrophotometer (Thermo-Fisher). Resulting data from growth curves was plotted with Graph Pad Prism 10.3.0 (GraphPad Software, LLC). SHI medium could not be used for growth curve assays as blood medium exhibited color changes and darkening due to 5% CO_2_ incubation. CFU/mL assays were used as alternative method for this medium. For CFU/mL assays in SHI medium, starters were prepared in SHI medium as outlined above, inoculated into 96 or 24 well plates. For each condition in triplicate, no peptide, 1 µM or 10 µM tryglysin A or revSHP (*S. mutans* reverse SHP peptide) were added to wells. Cultures were then incubated at 37°C with 5% CO_2_. Starting at time 0 (inoculation time after peptide addition), a portion of each sample was transferred to a separate 96 well plate, diluted for CFU/mL in TSB or sterile PBS, and on either TSA or TSA plus 5% defibrinated sheep blood in technical triplicate for CFU/mL. Plates were then incubated at 37°C with 5% CO_2_ for 1-2 days and colonies enumerated. Resulting CFU/mL for each condition was plotted using Graph Pad Prism 10.3.0, and examined for statistical significance between conditions at each time point via a One-way ANOVA with Tukey’s Multiple Comparisons Post-test.

### Growth curves, pH, and CFU/mL assays of salivary inoculum with tryglysin A

Norway collection saliva stocks at -80°C were thawed and inoculated into anaerobically incubated CDM at a concentration of 2 µL/mL CDM in sterile 15 mL tubes. For each condition in duplicate, 10 nM, 100 nM, 1 µM tryglysin A or *S. mitis* reverse SHP peptide (revSHP) or equivalent volume of PBS was added to cultures. For growth curves and pH measurements, cultures were grown in an anaerobic chamber at 37°C for 24 hours. Samples were taken at 0, 2, 4, 6, 8, 10 and 24 hours, and measured for growth via observation of OD_600_ and pH. Resulting growth curve data was plotted using Graph Pad Prism 10.3.0. Resulting pHs were plotted using Graph Pad Prism 10.3.0, and examined for statistical significance between conditions at each time point via a One-way ANOVA with Tukey’s Multiple Comparisons Post-test. For CFU/mL assays, samples were grown in the same manner with the exception that samples were taken at 0, 8, 12, and 24 hours for CFU/mL. Samples diluted for CFU/mL in sterile PBS at each time point and plated on either TSA or TSA plus 5% defibrinated sheep blood in technical triplicate for CFU/mL. Plates were then incubated at 37°C in the anaerobic chamber 24 to 48 hours and colonies enumerated. Resulting CFU/mL values were plotted using Graph Pad Prism 10.3.0, and examined for statistical significance between conditions at each time point via a One-way ANOVA with Tukey’s Multiple Comparisons Post-test.

### Growth and harvest of salivary samples with tryglysin A for sequencing

Norway collection saliva stocks at -80°C were thawed and inoculated into CDM as previously outlined in *Growth curves, pH, and CFU/mL assays of salivary inoculum with tryglysin A.* For each condition in duplicate, 1 µM tryglysin A, 1 µM revSHP, or equivalent volume of PBS was added to cultures. Samples were grown in an anaerobic chamber for 24 hours, at which time samples were harvested and then stored with 20% glycerol at -20°C. For direct sequencing of saliva, (i.e. initial inoculum sample), 1 mL of saliva was centrifuged at 14,000*xg* for 5 min. Supernatant was removed and remaining pellet was processed using the Masterpure Gram Positive DNA Purification Kit (Biosearch Technologies, MGP04100) according to manufacturer’s instructions. Samples and DNA (from bacteria and fungi from initial saliva) were then shipped on dry ice to the University of Illinois Chicago, under University of Illinois Chicago Institutional Review Board approval STUDY2023-0671, not human research determination; and CDC PHS Permit Number 20230524-2079A. Samples were thawed and additional sample DNA was extracted according to the manufacturer’s protocols using a Masterpure Gram Positive DNA Purification Kit (Biosearch Technologies, MGP04100). Sample purity was measured using a nanodrop (Thermo Fisher Nanodrop 2000 Spectrophotometer), samples acceptable for sequencing were those with an A28/A260 Of 1.8-2.0 and an A230/A260 of approximately 2.0. A minimum of 200 ng of total sample with a minimum concentration of 10 ng/µL were then submitted to the RUSH Genomics and Microbiome Core Facility (RUSH University) for shotgun metagenomics and ITS amplicon sequencing (ITS1-F to ITS2R, fungal ITS1 region). Sample data was processed as outlined in *Shotgun metagenomics sequencing and data processing* and *ITS and 16S amplicon sequencing and data processing*.

### Shotgun metagenomics sequencing and data processing

Shotgun metagenomic sequencing libraries were prepared with Illumina DNA Prep kit (Illumina, 20060059) then sequenced on NovaSeq X with 2x150 bp amplification. Resulting sequencing data for shotgun metagenomics was then submitted to the RUSH Bioinformatics Core (RUSH University). Raw data was checked using FastQC (https://www.bioinformatics.babraham.ac.uk/projects/fastqc/), followed by quality filtering and trimming using the algorithm bbduk (http://jgi.doe.gov/data-and-tools/bb-tools/). Short read taxonomic annotation was performed using the software package MetaPhlAn3 (v4.0.1)^37^ and functional gene annotation with HUMAnN3 (v3.5)^37^ mapping to the UniRef90 catalog (UniRef release 2019_01). UniRef90 relative abundance tables were regrouped into higher-level organizations, including: MetaCyc pathways, KEGG orthology, and UniProt gene families for downstream analysis. Resulting data, such as MetaPhlAn3 taxonomy assignments and BIOM files were imported into R Studio ver. 2024.09.0+375 (Posit Software, PBC) and data was processed using the R packages dplyr and tidyr^38,39^, to generate a stacked bar plot for relative abundances using the ggplot2 package^40^. PCoA plots were generated from MetaPhlAn taxonomic abundance data imported into R Studio and processed using the R packages metaphlanToPhyloseq, phlyoseq, and dplyr^39,41,42^. Ordination plots for PcoA were generated from taxonomic abundance data using the ordinate function in the phyloseq package with the PCoA method and Jaccard distance settings. PCoAs plots were plotted using ggplot2 and phyloseq packages in R Studio, with the imported MetaPhlAn3 taxonomic abundance data and ordination plots generated by the phyloseq package. Total read counts obtained from bbduk quality filtering and trimming were then used to estimate the relative count values based on MetaPhlAn3 taxonomic abundance data. These relative count values were used to perform statistical analysis of MetaPhlAn3 taxonomic abundance using DESeq2^25^. For analysis of shotgun metagenomics alpha diversity metrics, the relative count values based on MetaPhlAn3 taxonomic abundance data were analyzed with the estimate_richness function in the phyloseq package in R Studio. Resulting values from alpha diversity metrics were plotted using Graph Pad Prism 10.3.0, and examined for statistical significance between conditions using a via a Kruskal-Wallis pairwise test with a Benjamini & Hochberg correction^43^. Statistical significance of beta diversity metrics via PERMANOVA (adonis function)^44^ was determined using the vegan R package and pairwiseAdonis function^45^. We did note that for shotgun metagenomics samples grown in SHI medium, a large number of reads from *Ovis aries* were observed in these samples, which were filtered out during the shotgun metagenomics bioinformatics analysis. This correlates with the use of sheep blood during SHI medium culturing.

Before deposition of metagenomics sequence results to NCBI SRA, sequences were screened and filtered for potential human host sequences using Bowtie2^46^ with the human genome GRCh37 (NCBI RefSeq accession: GCF_000001405.13). Additional human read scrubbing was also performed by the Sequence Read Archive. All raw sequence reads for experiments are deposited to the NCBI SRA under BioProject PRJNA1189147.

### ITS and 16S amplicon sequencing and data processing

Genomic DNA was PCR amplified with primers sIDTP5_ITS1F and sIDTP7_ITS2R (<u>CTACACGACGCTCTTCCGATCT</u>CTTGGTCATTTAGAGGAAGTAA and <u>CAGACGTGTGCTCTTCCGATCT</u>GCTGCGTTCTTCATCGATGC respectively –underlined regions represent linker sequences). Amplicons were generated using a two-stage PCR amplification protocol similar to that described previously^47^. The primers contained 5’ common sequence tags, underlined (sIDTP5 and sIDTP7) that match 3’ sequences present in IDT xGen™ Amplicon UDI primers (part numbers: 10009846, 10009851, 10009852, and 10009853). First stage PCR amplifications were performed in 10 microliter reactions in 96-well plates, using repliQa HiFi ToughMix (Quantabio).

Genomic DNA input was 4 microliters per reaction. PCR conditions were 98°C for 2 minutes, followed by 28 cycles of 98°C for 10 min, 52°C for 1 min and 68°C for 1 min. Subsequently, a second PCR amplification was performed in 10 µL reactions in 96-well plates using repliQa HiFi ToughMix. Each well received a separate primer pair containing unique dual indices (i.e., from the IDT xGen™ amplicon UDI primer sets). One microliter of PCR product from the first stage amplification was used as template for the 2nd stage, without cleanup. Two microliters of primer were used per reaction. Cycling conditions were 98°C for 2 minutes, followed by 10 cycles of 98°C for 10 min, 60°C for 1 min and 68°C for 1 min.

The PCR product was pooled prior to a 0.6X Ampure cleanup, followed by size selection on the Pippin Prep device (Sage Science) with 1.5% agarose gel. Size selected DNA fragments (300-750 bp) from the PippinPrep were sequenced with a 10% phiX spike-in on an Illumina Miniseq sequencer employing a mid-output flow cell (2x154 paired-end reads). Based on the results of the MiniSeq sequencing run, samples were re-pooled to balance output evenly among samples. The pooled products were cleaned as described above (i.e., Pippin Prep and Ampure cleanups) and sequenced on an Illumina NovaSeq6000 instrument using a 500-cycle SP flow cell with read lengths extended to 2x259 bases. Library preparation, pooling, and MiniSeq sequencing were performed at the Genomics and Microbiome Core Facility (GMCF) at Rush University. NovaSeq sequencing was performed at the DNA Services Facility at the Roy J. Carver Biotechnology Center at the University of Illinois at Urbana-Champaign.

For 16S sequencing, genomic DNA was prepared and sequencing using the same method as for ITS sequencing, with the following changes. 16S sequences were PCR amplified with primers sIDTP_515F and sIDTP7_806R (<u>CTACACGACGCTCTTCCGATCT</u>GTGCCAGCMGCCGCGGTAA and <u>CAGACGTGTGCTCTTCCGATCT</u>GGACTACHVGGGTWTCTAAT – underlined regions represent linker sequences) respectively. For 16S sequencing, only the 0.6X Ampure cleanup was performed after the second round of PCR amplification before sequencing on the Illumina Miniseq sequencer, no PippinPrep size selection was performed. All other steps were the same as performed for ITS sequencing. Resulting sequencing data was then imported into QIIME 2 v.2024.5^48^ as demultiplexed sequences. Phred qualities of sequences were examined before proceeding with QIIME2 analysis, using a custom script. Sequences were trimmed using the q2-cutadapt trim-paired plugin with according primers using cutadapt^49^ followed by processing in one of the two manners: quality filtering via the q-score using q2-quality-filter-q-score plugin and then using Deblur^50^ (via q2-deblur) or alternatively using DADA2^51^ (via q2-dada2) to perform clustering, chimera filtering, and abundance filtering. The resulting table output from Deblur or DADA2 was used to select a set sampling depth for analysis of samples, which was set the same for all samples within an individual experiment. Selection of appropriate sample depth was determined via examination of alpha rarefaction values for the selected sampling depth using q2-diversity-alpha-rarefaction plugin. Phylogenetic trees were generated for diversity analysis using q2-phylogeny align-to-tree-mafft-fasttree using MAFFT and fasttree2^52,53^, which was then used to examine alpha and beta diversity metrics in QIIME 2. Alpha diversity metrics generated included Faith’s Phylogenetic Diversity, Chao1 index, observed features, Simpson’s Dominance, and Shannon entropy^54–57^. Beta diversity metrics and Principle Coordinate Analysis (PCoA) were estimated using q2-diversity, and included Jaccard and Bray-Curtis distance^58^. Alpha diversity statistical significance was determined via a Kruskal-Wallis pairwise test with a Benjamini & Hochberg correction^43^, while beta diversity statistical significance was determined using PERMANOVA (adonis function)^44^. Statistical significance was determined if q-value < 0.05. To determine genus presence within samples, taxonomy was assigned to ASVs using the q2-feature-classifier^59^ classifier-sklearn naïve Bayes taxonomy classifier or q2-feature-classifier classify-consensus-blast as follows: for 16S results against the Greengenes 2022.10 full length sequences^60^; for ITS results against the QIIME UNITE database v.10.0 (released 2024-04-04^61^). The QIIME UNITE database classifier was trained using the Nova High Performance computing cluster (HPC) at Iowa State University, while the QIIME Greengenes classifier was trained on a personal computer. Resulting classification bar plots, alpha, and beta diversity analysis were visualized using the QIIME2 View online platform^48^. PCoA plot results generated via QIIME 2 were imported into R Studio using the qiime2R package v0.99.6 (Bisanz, JE. GitHub) and plotted using ggplot2^40^.

Before deposition of 16S and ITS sequence results to NCBI SRA, sequences were screened and filtered for potential human host sequences using bowtie2^46^ on the Iowa State University Nova HPC with the GRCh37 human genome (NCBI RefSeq assembly: GCF_000001405.13). All raw sequence reads for experiments are deposited to the NCBI SRA under BioProject PRJNA1189147.

### Growth and harvest of salivary samples in planktonic or biofilm models with 5% CO_2_ for sequencing and CFU/mL analysis

Saliva samples from Chicago collection stored at -80°C were thawed and inoculated into SHI medium in the following manner. Each condition was grown in triplicate. For planktonic samples, 2 mL of salivary inoculum was added 3 mL of SHI medium in 15 mL conical tubes. For biofilm samples, a 1:100 dilution of saliva was added to SHI medium and aliquoted into 1 mL volumes into a 24 well plate. Samples were then grown at 37°C with 5% CO_2_ for 24 hours. At 0 hours (inoculation time) and 24 hours, a portion of each sample in triplicate was transferred to a separate 96 well plate, diluted for CFU/mL in TSB or sterile PBS, and plated on TSA plus 5% defibrinated sheep blood in technical triplicate for CFU/mL. Plates were then incubated at 37°C with 5% CO_2_ for 1-2 days, colonies enumerated, and plates imaged using a Genesys gel dock. Resulting CFU/mL for each condition was plotted using Graph Pad Prism 10.3.0, and examined for statistical significance between conditions at each time point via a One-way ANOVA with Tukey’s Multiple Comparisons Post-test. At 24 hours in the experiment, 1 mL each of cultures were also stored at -20 °C for downstream DNA extraction. Additionally, 1 mL of each culture in duplicate at 24 hours were stored with 20% glycerol at -80°C as “cultured consortia” for use in downstream experiments.

For DNA extraction, an initial saliva sample was sequenced as well as 1 mL each of experimental samples at 24 hours. For direct sequencing of saliva, (i.e. initial inoculum sample), 3 mL of saliva was centrifuged at 14,000*xg* for 5 minutes. Supernatant was removed and remaining pellet was processed using Masterpure Gram Positive DNA Purification Kit according to manufacturer’s instructions. Experimental samples were processed in the same manner. Sample purity for each was measured using a nanodrop (Thermo Fisher Nanodrop 2000 Spectrophotometer), samples acceptable for sequencing were those with an A28/A260 Of 1.8-2.0 and an A230/A260 of approximately 2.0. A minimum of 200 ng of total sample with a minimum concentration of 10 ng/µL were then submitted to the RUSH Genomics and Microbiome Core Facility (RUSH University) for shotgun metagenomics, ITS amplicon sequencing (ITS1-F to ITS2R, fungal ITS1 region), and 16S sequencing (515F and 806R, bacterial 16S rRNA region). Sample data was processed as outlined in *Shotgun metagenomics sequencing and data processing* and *ITS and 16S amplicon sequencing and data processing*.

### Growth and harvest of salivary samples or re-cultured consortium in CDM or SHI media with 5% CO_2_ for sequencing and CFU/mL analysis

Saliva samples from Chicago collection or “cultured consortia” stored at -70°C were thawed and inoculated into SHI medium or CDM in the following manner. Each condition was grown in triplicate. Samples were diluted 1:100 into 6 mL of SHI or CDM, and then incubated for 24 hours at 37°C with 5% CO_2_. At 0 hours and 24 hours, initial inoculum or cultures were harvested, plated for CFU/mL, and results analyzed as outlined in *Growth and harvest of salivary samples in planktonic or biofilm models with 5% CO2 for sequencing and CFU/mL analysis.* At 24 hours in the experiment, 1 mL each of cultures were also stored at -20 °C for downstream DNA extraction.

For DNA extraction, 1 mL of each sample at 24 hours was centrifuged at 14,000*xg* for 5 minutes. Supernatant was removed and remaining pellet was processed using Masterpure Gram Positive DNA Purification Kit according to manufacturer’s instructions. Sample purity for each was measured using a nanodrop as previously mentioned. A minimum of 200 ng of total sample with a minimum concentration of 10 ng/µL were then submitted to the RUSH Genomics and Microbiome Core Facility (RUSH University) for ITS amplicon sequencing and 16S sequencing as previously described. Sample data was processed as outlined in *Shotgun metagenomics sequencing and data processing* and *ITS and 16S amplicon sequencing and data processing*.

## DATA AVAILABILITY

All shotgun metagenomics data, ITS, and 16S data has been deposited to the NCBI Sequence Read Archive (SRA) under BioProject PRJNA1189147.

## CODE AVAILABILITY

All codes and packages are available via their respective citations in *Materials and Methods*. The custom Linux shell script used to examine Phred score is available on GitHub (https://github.com/b-rued/Phred-check).

## ACKNOWLEDGEMENTS

The authors would like to thank the members of the Federle and Petersen labs for helpful discussions and proofreading of the manuscript. Additionally, we specifically acknowledge Heidi A. Åmdal from the Petersen lab for her assistance with experimental procedures, analysis, and insight on findings. This study was supported by the National Institutes of Health (R01-AI162679 to M.J.F., K99DE032311 and R00DE032311 to B.E.R.). Additional collaboration and networking activities were supported by the Research Council of Norway (RESISFORCE, Grant 322375 to F.C.P). Library preparation, pooling, and sequencing were performed at the Genomics and Microbiome Core Facility (GMCF - https://tinyurl.com/bdz6j64f) at Rush University. Shotgun metagenomics short-read annotation and bioinformatics analysis were performed at the Rush Research Bioinformatics Core (RRBC - https://tinyurl.com/8mjxs55d) at Rush University. The research reported in this paper is also partially supported by the HPC@ISU equipment at Iowa State University, some of which has been purchased through funding provided by NSF under MRI grants number 1726447 and MRI2018594.

## AUTHOR CONTRIBUTIONS

Conceptualization, B.E.R, M.J.F, and F.C.P.; methodology, B.E.R., D.A., R.P.P., F.C.P.; resources, B.E.R, M.J.F., F.C.P., R.P.P., B.C.C., and M.R.S.; data collection and analysis, B.E.R, D.A., M.J.F, and F.C.P.; writing – original draft preparation, B.E.R.; writing – review and editing, B.E.R, D.A., M.J.F., F.C.P., R.P.P., B.C.C., and M.R.S.; project administration and oversight, B.E.R, M.J.F., and F.C.P..; funding acquisition, B.E.R, M.J.F., and F.C.P. All authors have read and agreed to the published version of the manuscript.

