## Supplemental Figures for "Impact of the RaS-RiPP tryglysin and culturing conditions on *ex-vivo* oral microbiomes"

[b]Institute of Oral Biology. University of Oslo, Oslo, Norway.

[c]Department of Chemistry. Princeton University, Princeton, New Jersey, USA.

[d]College of Dentistry. University of Illinois - Chicago, Illinois, USA.

[e]Department of Pharmaceutical Sciences. University of Illinois - Chicago, Chicago, Illinois, USA

### SUPPLEMENTAL FIGURES

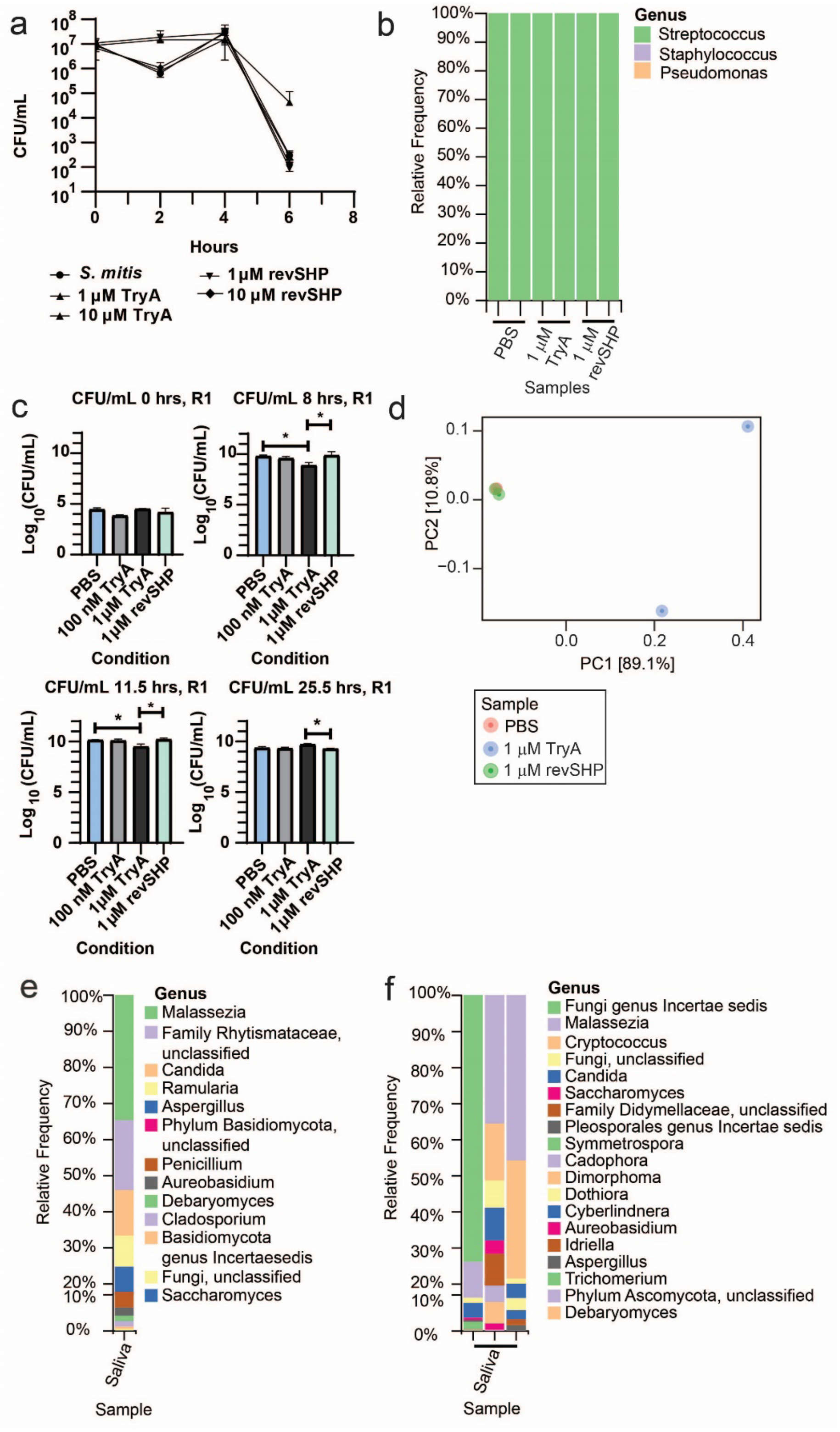

**FIGURE S1:** Additional data on initial salivary inoculum and examining differences in Tryglysin A inhibition between SHI medium and CDM. A) CFU/mL of wild-type *S. mitis* from Chicago laboratory exposed to increasing TryA in SHI. Concentrations of Tryglysin A are indicated in the legend below the graph and correspond to the respective symbol types. This experiment was performed three times with similar results. B) 16S rRNA sequencing results of samples grown in CDM for 24 hours, with PBS, 1 µM Tryglysin A, or 1 µM revSHP (*S. mutans* reverse SHP, Table 1). C) CFU/mL bar graphs of salivary inoculum in CDM medium exposed to increasing TryA, PBS, or revSHP. This experiment was performed twice. Data correlates to graph shown in Fig. 1E. D) Jaccard PCoA plot of metagenomics results from the same samples as presented in Fig. 1F. E) Fungal phyla detected in human pooled saliva isolates as determined by ITS sequencing from Norway isolation. Relative abundance is shown for each sample, phyla are indicated by color as detailed in the legends beside each graph. The Norwegian pooled saliva was sequenced once. F) Fungal phyla detected in human pooled saliva isolates as determined by ITS sequencing from Chicago isolation. Relative abundance is shown for each sample, phyla are indicated by color as detailed in the legends beside each graph. The Chicago pooled saliva was sequenced in triplicate, with all samples originating from DNA extracted from the same salivary pool. Note that this data is also presented alongside additional samples in Fig. S2B.

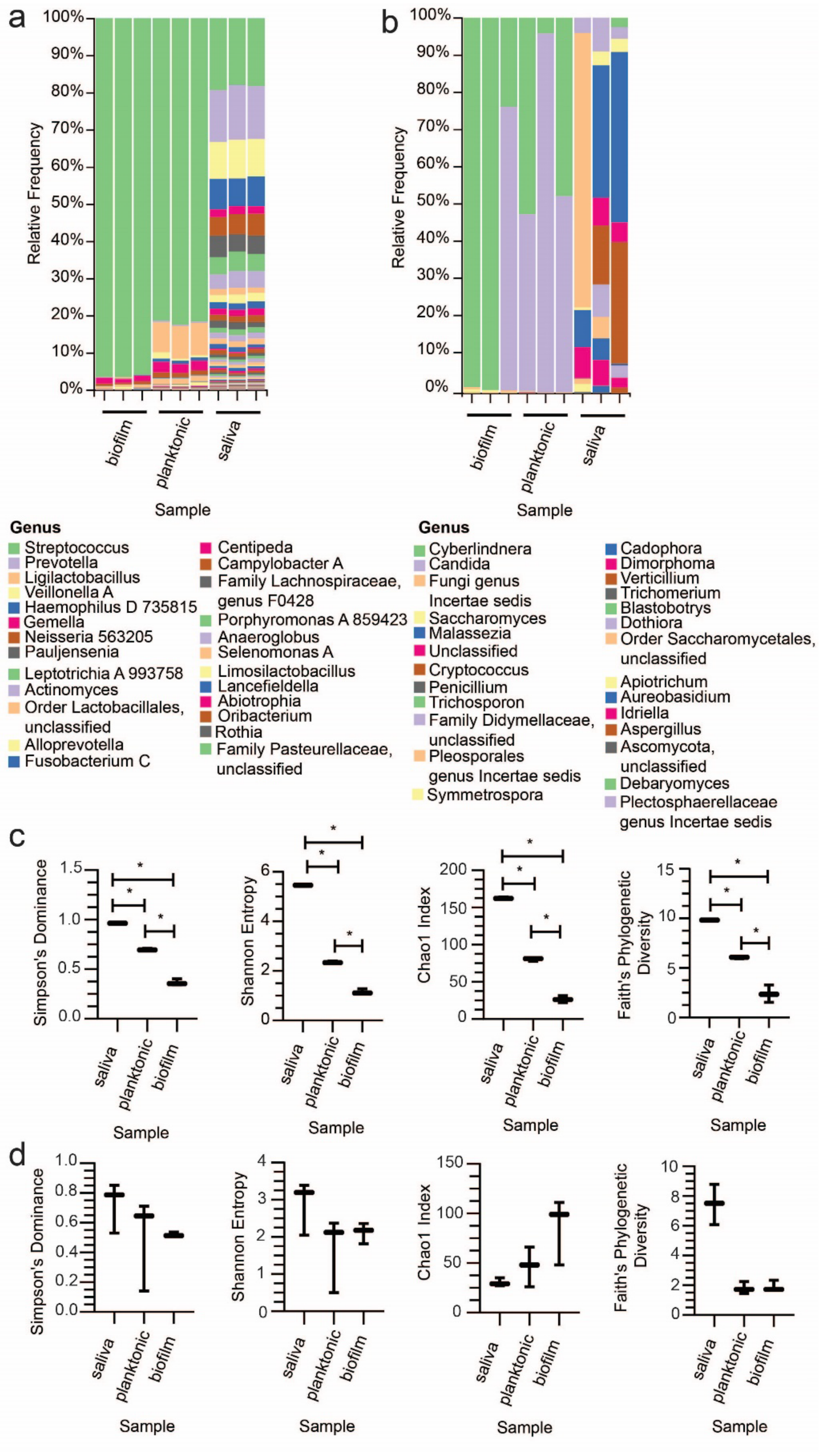
a
b
Relative Frequency
100%
90%
80%
70%
60%
50%
40%
30%
20%
10%
0%
biofilm
planktonic
saliva
Sample
Genus
Streptococcus
Prevotella
Ligilactobacillus
Veillonella A
Haemophilus D 735815
Gemella
Neisseria 563205
Pauljensenia
Leptotrichia A 993758
Actinomyces
Order Lactobacillales, unclassified
Alloprevotella
Fusobacterium C
Centipeda
Campylobacter A
Family Lachnospiraceae, genus F0428
Porphyromonas A 859423
Anaeroglobus
Selenomonas A
Limosilactobacillus
Lancefieldella
Abiotrophia
Oribacterium
Rothia
Family Pasteurellaceae, unclassified
Genus
Cyberlindnera
Candida
Fungi genus Incertae sedis
Saccharomyces
Malassezia
Unclassified
Cryptococcus
Penicillium
Trichosporon
Family Didymellaceae, unclassified
Pleosporales genus Incertae sedis
Symmetrospora
Cadophora
Dimorphoma
Verticillium
Trichomerium
Blastobotrys
Dothiora
Order Saccharomycetales, unclassified
Apiotrichum
Aureobasidium
Idriella
Aspergillus
Ascomycota, unclassified
Debaryomyces
Plectosphaerellaceae genus Incertae sedis
c
Simpson's Dominance
Shannon Entropy
Chao1 Index
Faith's Phylogenetic Diversity
saliva
planktonic
biofilm
Sample
d
Simpson's Dominance
Shannon Entropy
Chao1 Index
Faith's Phylogenetic Diversity
saliva
planktonic
biofilm
Sample

**FIGURE S2:** Additional data examining the use of 5% $CO_2$ as a culturing method for *ex-vivo* oral microbiomes. All samples shown were grown in SHI media for 24 hours with 5% $CO_2$. For panels C and D, statistical significance between conditions was determined using a pairwise Kruskal-Wallis test with a p-value adjustment using a Benjamini & Hochberg correction. * indicates q-value < 0.05. If no symbols are present, this indicates comparisons were non-significant. A) 16S sequencing results showing bacterial genera detected in a biofilm or planktonic model of growth, compared to genera detected in the original salivary inoculum. The top 25 genera are indicated by the figure legend, for all genera detected see Table 9 in Supplementary File 02. B) ITS sequencing results showing fungal genera detected in a biofilm or planktonic model of growth, compared to genera detected in the original salivary inoculum. For genera detected in each sample see Table 7 in Supplementary File 02. C) Alpha diversity metrics for 16S sequencing results shown in panel A. D) Alpha diversity metrics for the ITS sequencing shown in panel B.

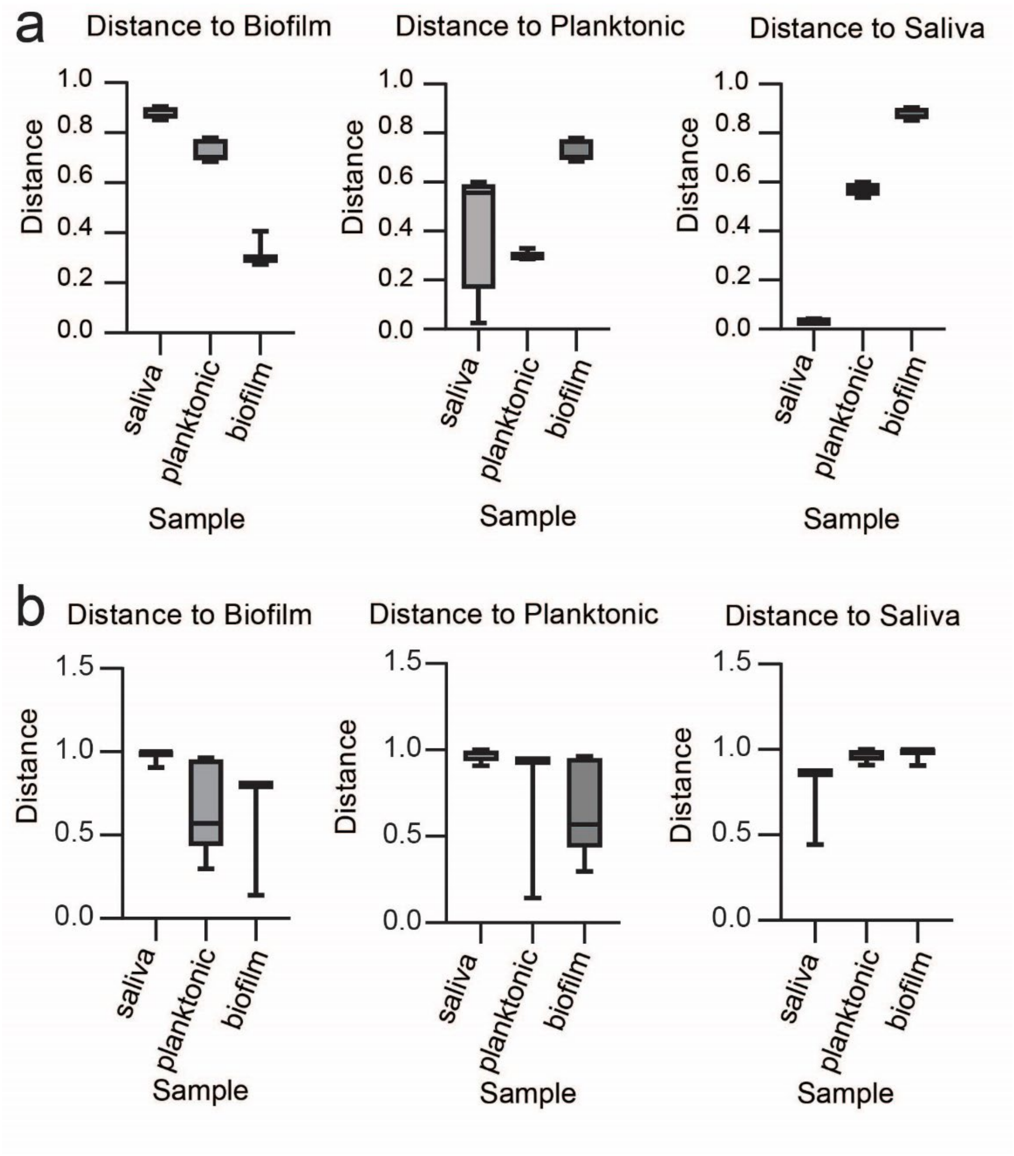


**FIGURE S3:** Beta diversity metrics for 16S and ITS sequencing from *ex-vivo* oral samples cultured with 5% $CO_2$. Statistical significance between conditions was determined using PERMANOVA (adonis function). Pairwise comparisons are indicated between samples by brackets. If no brackets are present, this indicates comparisons were non-significant. A) Beta diversity metrics determined via Jaccard distance for 16S sequencing results shown in Fig. S2A. B) Beta diversity metrics determined via Bray-Curtis distance for ITS sequencing results shown in Fig. 3B and Fig. S2B.

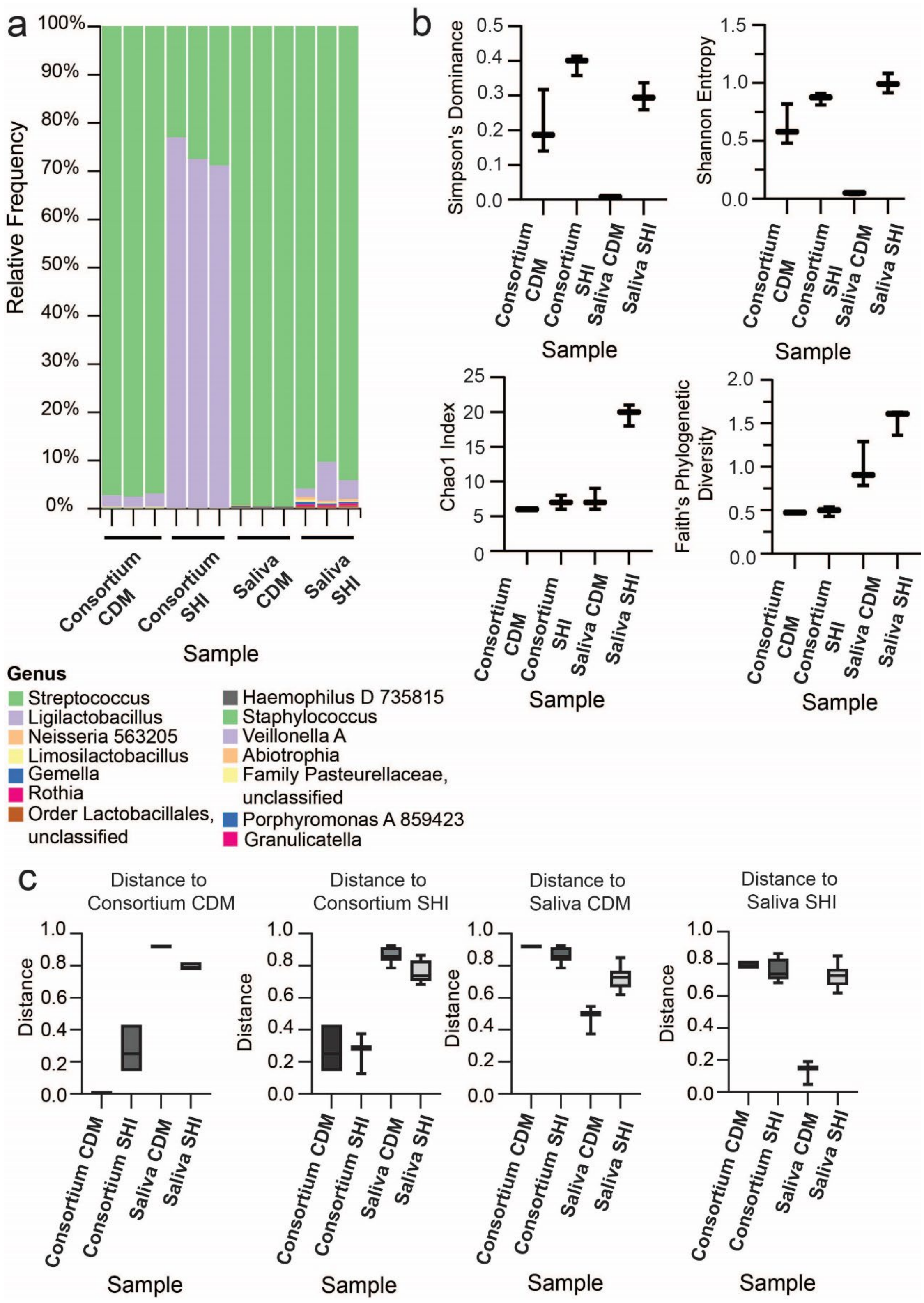


**FIGURE S4:** Additional data examining the impact of CDM vs. SHI media on bacterial population for culturing *ex-vivo* oral microbiomes. Samples were grown in CDM or SHI media for 24 hours with 5% $CO_2$, in biological triplicate. For graphs in panels B and C, note the differences in scales on the y-axis. A) 16S sequencing results showing bacterial genera detected after growth of saliva inoculum or previously stored consortia in CDM or SHI media. B) Alpha diversity metrics with samples split by inoculum source and media for 16S sequencing results shown in panel A. Statistical significance between conditions was determined using a pairwise Kruskal-Wallis test with a p-value adjustment using a Benjamini & Hochberg correction. If no symbols are present, this indicates comparisons were non-significant. C) Beta diversity metrics using Jaccard distance with samples split by inoculum source and media for 16S sequencing results shown in panel A. Statistical significance between conditions was determined using pairwise adonis PERMANOVA (adonis function). Pairwise comparisons are indicated between samples by brackets. If no brackets are present, this indicates comparisons were non-significant.

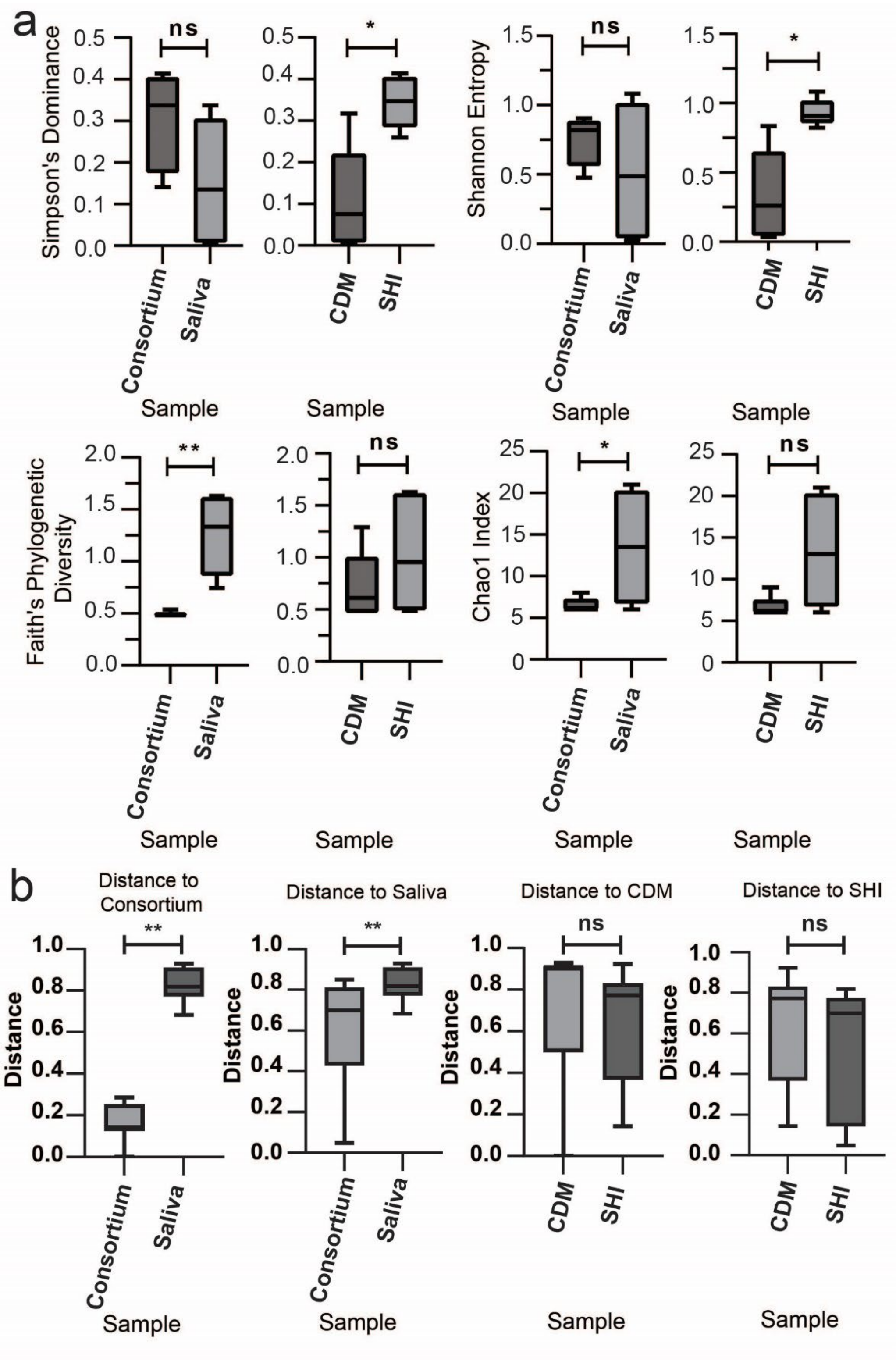


**FIGURE S5:** Comparison of additional alpha and beta diversity metrics for 16S data shown in Figure 4 and S4. Shown are two other comparisons: consortia vs. saliva (inoculum source), or CDM vs. SHI media (medium condition). For all graphs, note the differences in scales on the y-axis. A) Alpha diversity metrics for additional comparisons. Statistical significance between conditions was determined using a pairwise Kruskal-Wallis test with a p-value adjustment using a Benjamini & Hochberg correction. *, q-value < 0.05; **, q-value < 0.005; ns, nonsignificant. B) Beta diversity metrics using Jaccard distance for additional comparison. Statistical significance between conditions was determined using pairwise PERMANOVA (adonis function). **, q-value < 0.005; ns, nonsignificant.

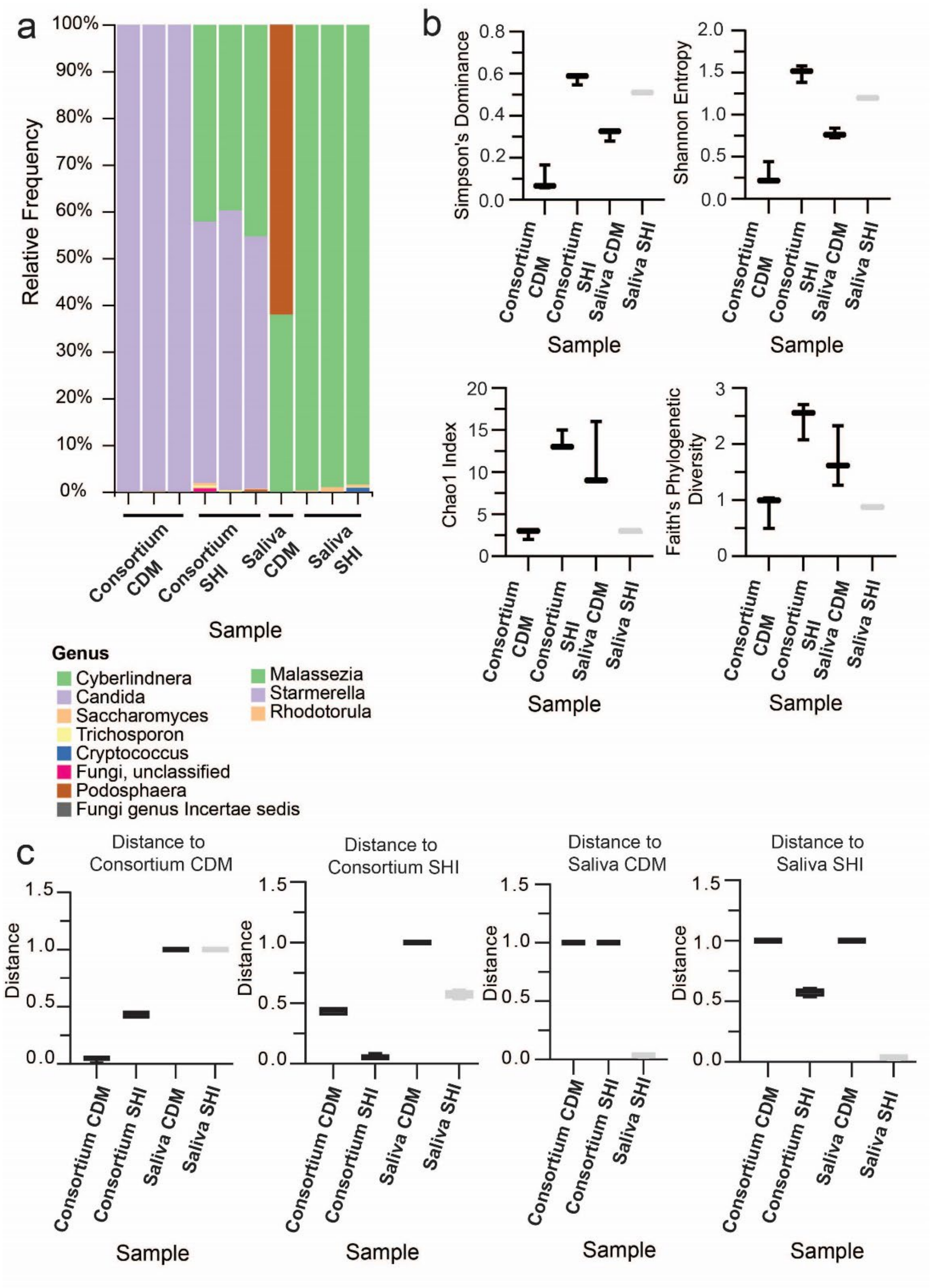


**FIGURE S6**: Additional data examining the impact of CDM vs. SHI media on fungal population for culturing *ex-vivo* oral microbiomes. Samples were grown in CDM or SHI media for 24 hours with 5% $CO_2$, in biological triplicate. For graphs in panels B and C, note the differences in scales on the y-axis. A) ITS sequencing results showing fungal genera detected after growth of saliva inoculum or previously stored consortia in CDM or SHI media. B) Alpha diversity metrics with samples split by inoculum source and media for ITS sequencing results shown in panel A. Statistical significance between conditions was determined using a pairwise Kruskal-Wallis test with a p-value adjustment using a Benjamini & Hochberg correction. If no symbols are present, this indicates comparisons were non-significant. C) Beta diversity metrics using Bray-Curtis distance with samples split by inoculum source and media for ITS sequencing results shown in panel A. Statistical significance between conditions was determined using PERMANOVA (adonis function). Pairwise comparisons are indicated between samples by brackets. If no brackets are present, this indicates comparisons were non-significant.

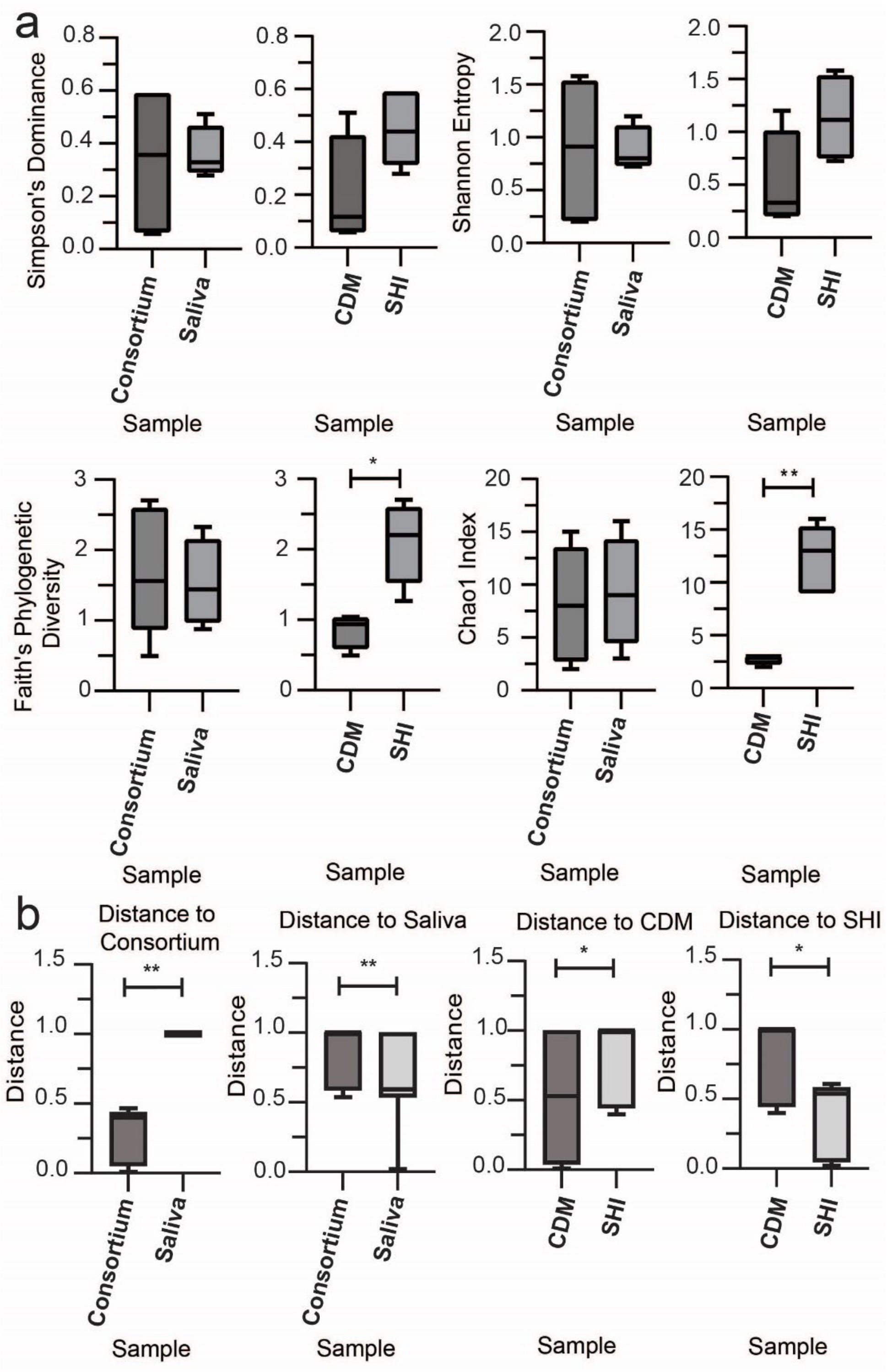


**FIGURE S7:** Comparison of additional alpha and beta diversity metrics for ITS data shown in Figure 4 and S6. Shown are two other comparisons: consortia vs. saliva, or

CDM vs. SHI media. For all graphs, note the differences in scales on the y-axis. A) Alpha diversity metrics for additional comparisons. Statistical significance between conditions was determined using a pairwise Kruskal-Wallis test with a p-value adjustment using a Benjamini & Hochberg correction. *, q-value < 0.05; **, q-value < 0.005; no symbols present, nonsignificant. B) Beta diversity metrics using Bray-Curtis distance for additional comparisons. Statistical significance between conditions was determined using pairwise PERMANOVA (adonis function). *, q-value < 0.05; no symbols present, nonsignificant.
