## Supplementary Data File 1 for "Impact of the RaS-RiPP tryglysin and culturing conditions on *ex-vivo* oral microbiomes"

[b]Institute of Oral Biology. University of Oslo, Oslo, Norway.

[c]Department of Chemistry. Princeton University, Princeton, New Jersey, USA.

[d]College of Dentistry. University of Illinois - Chicago, Illinois, USA.

[e]Department of Pharmaceutical Sciences. University of Illinois - Chicago, Chicago, Illinois, USA

**TABLE S1.** Bacterial strains and primers used in this study.[a]

| Bacterial strains | | | |
|---|---|---|---|
| **Strain** | **Description** | **Antibiotic Resistance** | **Reference[b]** |
| *S. mitis* | *Streptococcus mitis* CCUG 31611 wild-type isolate, also known as NCTC 12261 | None | Carlsson 1968. *Odontol Revy.* |
| *S. ferus* | *Streptococcus ferus* DSM20646 | None | Bushin *et al.*, 2018. *JACS.* |
| **Primers used for 16S or ITS sequencing** | | | |
| **Primer** | **Sequence (5' to 3')** | **Sequencing Target** | **Reference[b]** |
| **515F** | GTGCCAGCMGCCGCGGTAA | Bacterial 16S rRNA V4 region | Caparoso *et al.*, 2010. *PNAS.* |
| **806R** | GGACTACHVGGGTWTCTAAT | Bacterial 16S rRNA V4 region | Caparoso *et al.*, 2010. *PNAS.* |
| **ITS1-F** | CTTGGTCATTTAGAGGAAGTAA | Fungal ITS1 region | Gardes and Bruns. 1993. *Mol. Eco.* |
| **ITS2R** | GCTGCGTTCTTCATCGATGC | Fungal ITS1 region | White *et al.*, 1990. |

[a]Strains were used as described in *Materials and Methods*.
[b]See attached reference list.
